# Spatiotemporal graph neural networks reveal conformational binding signature in protein dynamics

**DOI:** 10.64898/2026.05.19.726195

**Authors:** Stefano Motta, Gianluca Santini, Samman Mansoor, Ferdoos Hossein Nezhad, Massimiliano Meli, Alessandro Pandini

**Author notes:** To whom correspondence should be sent,.

## Abstract

Biomolecular function is often controlled by structural and dynamical adaptations to binding events. Although molecular dynamics (MD) simulations can capture these events at atomic resolution, separating functional signatures from stochastic noise remains challenging. Traditional methods often struggle to isolate mechanistically relevant differences across independent replicas. Here, we introduce an explainable deep learning approach that learns state-specific dynamic signatures directly from MD trajectories. By coupling a dynamic protein graph representation with group-aware contrastive learning across independent replicas, the model detects the signatures, filtering out trajectory-specific correlations. An explainable AI framework then maps the identified differences on individual residues. We demonstrate this approach by identifying “binding-ready” conformations in a T4-Lysozyme mutant, recovering the allosteric determinants of peptide recognition in the PDZ3 domain, and isolating a ligand-independent activation signature for the A2A receptor. Our GISTnet-MD method generalizes across unseen data during comparative MD analysis, translating raw trajectory differences into residue-level determinants of protein function.

## Introduction

Protein functions are often mediated by conformational dynamics, which can be driven by structural adaptations to binding events^1^. The interaction with a physiological or a pharmacological molecule acts as a thermodynamic perturbation, shifting the conformational equilibrium and, in some cases, enabling allosteric responses^2,3^. Quantifying these dynamic rearrangements is important to characterize the effect of pathogenic mutations and to inform the design of drugs and allosteric modulators, applications that increasingly rely on the integration of machine learning and graph theory^4,5^. Molecular Dynamics (MD) simulations are traditionally used to generate atomistic descriptions of these conformational events. In this context, a major challenge for computational biophysics is comparative MD analysis: the comparison of trajectories generated from distinct states (e.g. apo versus holo, or wild-type versus mutant, etc.) to identify their functional features. Indeed, one of the main goals in this field has been to extract the structural and dynamical features that invariantly define a specific state across independent replicas (i.e. simulations initiated from the same coordinates but with different randomized initial velocities to ensure independent sampling)^6,7^. However, distinguishing the thermodynamic signatures of functional binding from stochastic thermal fluctuations and replica-specific variance is still challenging.

Established methods to detect and annotate conformational states rely on geometric measurements (e.g., RMSD, RMSF), dynamic cross-correlation matrices (DCCM), or dimensionality reduction techniques such as Principal Component Analysis (PCA) and time-lagged independent component analysis (tICA)^8^. To explicitly compare structural dynamics across distinct conformational states, specialized classical algorithms have been proposed, such as the evaluation of differential inter-residue contact maps^9^ or the application of mutual information metrics to capture anharmonic spatial correlations^10^. In the same context, techniques such as Iterative Linear Discriminant Analysis (LDA-Iter)^11^ have been applied to directly compare distinct trajectories. Because these methods rely on linear projections, structural alignments, or average covariance matrices, the dynamic signatures are often masked by baseline thermal fluctuations.

More recently, Deep Learning (DL) architectures were introduced to address the limits of linear models. Frameworks based on the Variational Approach for Markov Processes (VAMP), such as VAMPnets^12^, substitute multi-step state definition pipelines with end-to-end differentiable networks that optimize slow kinetic modes. In parallel, methods such as DeepTICA^13^ and the State Predictive Information Bottleneck (SPIB)^14^ utilize non-linear Koopman operators and information theory to construct low-dimensional reaction coordinates that preserve mutual information regarding future metastable states. Furthermore, architectures such as Deep Targeted Discriminant Analysis (Deep-TDA)^15^ extend classical discriminant frameworks by employing neural networks to project structural descriptors from multiple metastable states onto separated Gaussian distributions, establishing a non-linear, data-driven reaction coordinate for state comparison. For the specific task of comparative analysis across protein variants, architectures like DiffNets^16^ employ self-supervised autoencoders to identify low-dimensional structural features that predict biochemical differences.

Advances in geometric deep learning have stimulated the use of Graph Neural Networks (GNNs) for the analysis of MD trajectories.^17^ By representing biomolecules as dynamic topological graphs, where nodes correspond to residues and edges encode spatial distances, GNNs are inherently invariant to 3D rotations and translations making them well suited to embed conformational ensemble^18^. This structural representation reduces the dependency on manually engineered collective variables, allowing the model to directly map both local chemical environments and global inter-residue communications. ^17,19,20^. Recent architectures, such as AlloPool^21^, extend this paradigm by employing temporal attention and graph aggregation to iteratively prune residue interactions, identifying time-dependent minimal networks governing allosteric responses.

Applying GNNs to the comparative MD analysis has specific methodological challenges. First, MD simulations generate temporally correlated and sparse data points.^22–24^ Consequently, end-to-end classification models are susceptible to overfitting: the network models learn decision boundaries based on stochastic thermal noise or trajectory-specific initial coordinates rather than generalized biophysical features that remain invariant across independent replicas^23,24^. This tendency is amplified when analyses are restricted to single simulation replicas. In these cases, isolating the functional signal from baseline structural fluctuations is particularly challenging.^25^ Second, neural network architectures generally lack inherent interpretability. To effectively support structural biology or drug design, GNN models cannot act as black boxes: the network must translate its mathematical classifications into understandable physical features, such as specific 3D interaction networks and functional residues. ^26^. Third, comparative MD analysis is frequently confounded by initial coordinate biases. When ensembles originate from structurally different crystal structures, neural networks tend to classify states based on the signature of the initial difference rather than the differences in functional dynamics.^22,27,28^

To address the aforementioned limitations, here we introduce GISTnet-MD (Graph Invariant SpatioTemporal Network for Molecular Dynamics). GISTnet-MD is an explainable deep learning framework designed to extract invariant dynamic signatures (i.e. underlying biophysical patterns that consistently define a structural ensemble across independent replicas) associated with binding events or structural perturbations. The architecture couples a spatial Graph Neural Network (GNN) with a temporal 1-Dimensional Convolutional Neural Network (1D-CNN). This enables the model to map protein conformations directly via a continuous graph representation while simultaneously capturing time-dependent dynamics from sliding trajectory windows. The predictive task is formulated as a classification problem: every temporal window provided from the input is assigned a target label corresponding to the biological condition of its parent simulation (e.g., “apo” versus “holo”, or “wild-type” versus “mutant”). By tasking the neural network to predict these predefined target label from the trajectory data, the model is directed to identify the underlying structural and dynamic features that consistently differentiate the chosen target ensembles. To separate functional events from stochastic thermal noise, the spatiotemporal module is trained using a group-aware contrastive learning protocol, which constrains the model to learn functional dynamic patterns rather than memorizing trajectory-specific noise. Following this representation learning phase, a linear classifier is trained to predict the assigned state labels from the optimized latent embeddings.

To validate the identified state-defining features, our approach uses a Leave-One-Replica-Out Cross-Validation (LORO-CV) protocol. Predictive performance is evaluated exclusively on hold-out trajectories, ensuring that the learned representations generalize across independent sampling events. Finally, to map the dynamics patterns back to specific three-dimensional protein structures, the framework integrates an Explainable AI (xAI) pipeline based on the Integrated Gradients (IG) technique. This approach provides interpretability by attributing importance scores to individual input features, effectively identifying the specific residues and interactions that drive the model’s predictions.

The scope of application of GISTnet-MD is evaluated across three molecular systems characterized by distinct mechanisms of structural adaptation to the binding event. ***T4-Lysozyme (Binding-ready cavity formation):*** As a highly controlled test case, we compared the dynamics of the wild-type (WT) protein against the L99A mutant. This canonical mutation creates an internal hydrophobic cavity, shifting the ensemble into a “binding-ready” state^29–31^. ***PDZ3 Domain (Dynamic allostery and peptide recognition):*** PDZ domains bind short C-terminal peptides in a reversible and highly dynamic manner. To evaluate the detection of distributed allosteric networks, we compared the apo and peptide-bound (holo) states. In this system, allosteric communication does not manifest as large-scale structural rearrangements, but rather through the propagation of subtle variations in thermal fluctuations, often termed “dynamic allostery”, along networks of non-covalent interactions^32–35^. ***Adenosine A2A Receptor (Generalized agonist signature):*** The Adenosine A2A receptor is a Class A G-protein-coupled receptor (GPCR) whose functional activation relies on transmitting allosteric signals from an orthosteric binding pocket to intracellular microswitches, ultimately coordinating macroscopic structural shifts^36–39^. To extract dynamic activation signatures, we evaluated the model on two comparative tasks: classifying agonist-bound versus antagonist-bound ensembles and comparing agonist-bound versus active-apo trajectories.

While the three selected systems are experimentally well-characterized, realistic predictive applications often lack experimental structures for perturbed states, such as a novel ligand-bound complex or an uncharacterized mutant. To emulate a realistic scenario, and to prevent the network from classifying states based on initial geometry biases, simulation data for comparative analyses (except the agonist vs. antagonist comparison for the A2A receptor) were generated from identical starting coordinates. Physically, the initial systems differed only by the explicitly defined perturbation, such as the in-silico introduction of a point mutation, a peptide, or a specific ligand. However, to avoid label-leakage during model training, the source of the perturbation was completely masked from the input graph. While the common starting structure guarantees that the response arises only from differences in relaxation dynamics, detecting the dynamical signature of this response requires a specialized deep learning architecture. By coupling a dynamic spatiotemporal GNN with a group-aware validation protocol, GISTnet-MD explicitly decouples functional events from stochastic fluctuations. Crucially, through a cross-validation protocol across independent replicas, the proposed training strategy avoids that importance is incorrectly assigned to thermal noise instead of signal in the training data. This ensures that the extracted IG scores highlight perturbation-related residues and interactions that generalize across unseen trajectories. Ultimately, the developed pipeline establishes a robust approach to successfully map the biophysical determinants of protein function directly onto residue networks.

## Results

### GISTnet-MD Architecture

To detect dynamic signatures defining distinct conformational ensembles, we developed the GISTnet-MD architecture (**Fig. 1A**). Since conformational differences in MD trajectories are often masked by stochastic noise, GISTnet-MD is designed to extract features that are invariant to the initial conditions and isolate the functional signal. Analyzing raw atomic coordinates typically requires prior structural alignment, a process that can introduce reference biases. To overcome this limitation, the model first represents the protein as a continuous spatial graph, where nodes correspond to amino acid residues and edges describe their geometrical proximity (**Fig. 1B**). This graph-based representation maintains translational and rotational invariance, eliminating the need for prior structural alignment. To provide more accurate description, inter-residue distances are expanded using radial basis functions (RBFs). Additionally, a continuous envelope function is applied to prevent numerical discontinuities when inter-residue distances fluctuate around the interaction cutoff. This combined approach provides a structural representation of the local chemical environment, ensuring that small fluctuations in graph connectivity do not introduce artifacts^40^.

**Fig. 1.**
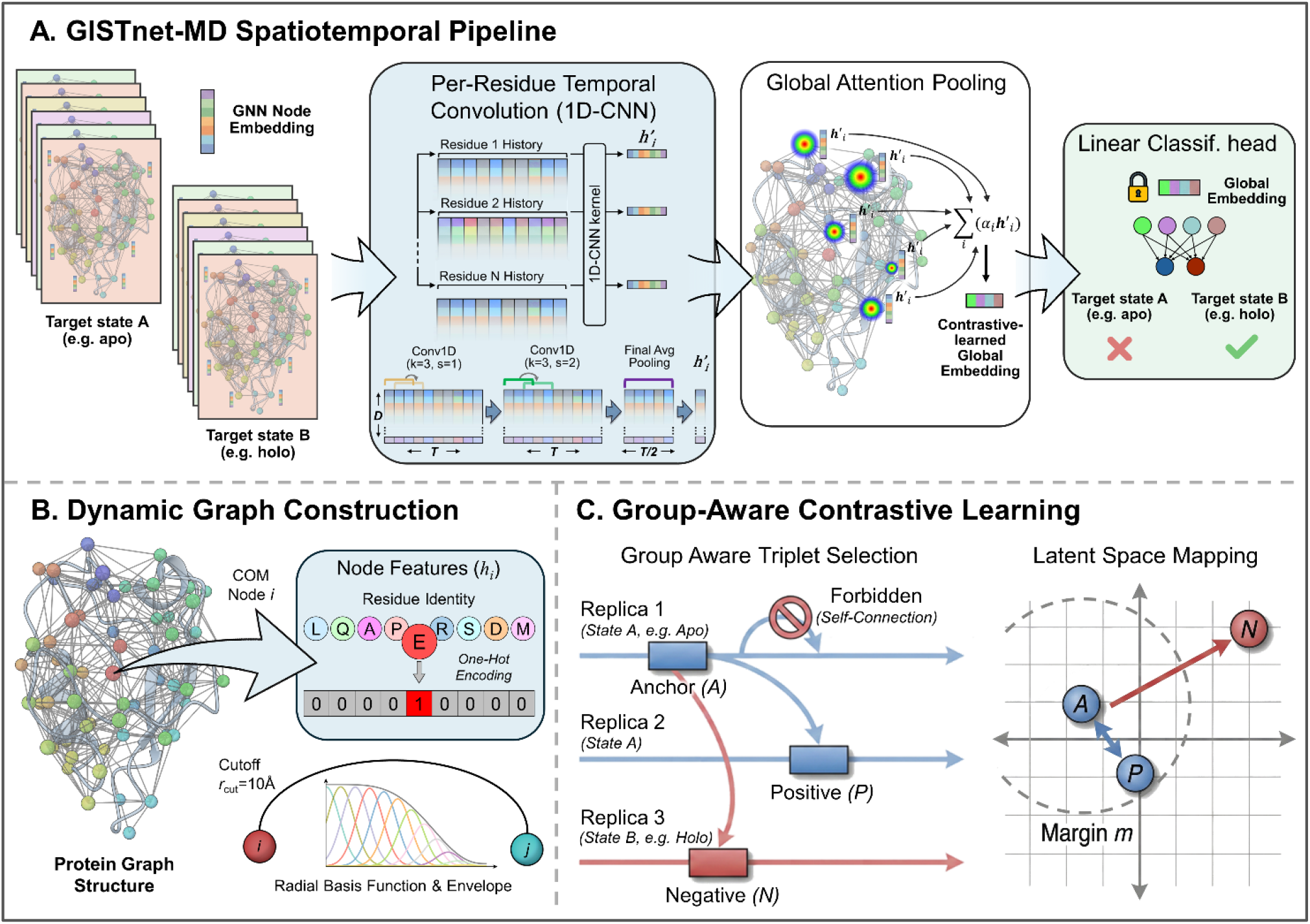
The GISTnet-MD architecture. **A: Spatiotemporal Pipeline**. A target label corresponding to the biological condition of the parent trajectory is assigned to each sliding temporal windows extracted from MD simulations (e.g., apo versus holo). Within each window, individual frames are represented as geometrical graphs to learn structural node embeddings. A per-residue temporal convolution (1D-CNN) slides independently over the temporal history of each residue, extracting local dynamic signatures and filtering high-frequency noise. A Global Attention Pooling mechanism assigns a weight α_i_ to each residue, compressing the temporal window into a single global embedding optimized via contrastive learning (detailed in C). Once optimized, the spatiotemporal backbone is frozen, and a linear classification head is trained on the fixed global embeddings to predict the functional target state. **B: Dynamic Graph Construction**. Details of the initial structural representation fed to the pipeline in ***A***. 3D coordinates from MD trajectories are converted into translationally and rotationally invariant graphs. Nodes represent amino acid residues initialized with one-hot identity vectors, while edges define spatial connectivity within a physical cutoff distance of 10 Å. Inter-residue distances are expanded via continuous Radial Basis Functions (RBF) and a cutoff envelope. **C: Group-Aware Contrastive Learning**. Details of the triplet selection strategy (left) and the resulting latent space mapping (right). The algorithm selects triplets consisting of an Anchor window (*A*), a Positive window (*P*) sampled from the same target state but from a different independent replica, and a Negative window (*N*) from the opposing target state; self-connections within the same simulation replica are strictly forbidden. As illustrated in the latent space mapping, the contrastive loss minimizes the Euclidean distance between *A* and *P* while maximizing the distance between *A* and *N*, enforcing that the negative sample is pushed outside a predefined geometric margin *m* relative to the Anchor.

Because target states (defined in our framework as functional states corresponding to specific simulation conditions such as apo/holo, or WT/mutant) are characterized by conformational ensembles rather than static snapshots, the model subsequently analyzes sliding temporal windows. A 1D-CNN processes the temporal sequence of the structural embeddings for each residue (**Fig. 1A**), acting as a dynamic filter that attenuates high-frequency thermal noise. A global attention pooling mechanism then evaluates these per-residue embedding vector, assigning an importance weight to each residue. This step compresses the temporal window into a single latent vector that encodes the target state of the protein. The necessity of temporal integration is supported by our ablation study, evaluated under a strict LORO-CV regime (**Table 1, Fig. 2**). When the temporal dimension is removed (Static End-to-End model), classification accuracy on unseen replicas drops significantly, indicating that functional states are driven by local dynamics rather than static conformational differences. Temporal shuffling experiments further confirmed that the 1D-CNN captures state-specific ensemble dynamics rather than memorizing exact temporal correlations (Supplementary Note 1).

**Table 1.** Architectural components of the evaluated model variants. The ablation matrix details the inclusion (✓) or exclusion (-) of key computational paradigms (temporal integration via 1D-CNN, multi-replica training, contrastive pre-training, and the group-aware constraint) across the tested deep learning frameworks. The corresponding generalization performances for each configuration are reported in **Fig. 2**.

|  | Temporal Integration<br>(1D-CNN) | Multi-Replica<br>Training | Contrastive<br>Pre-training | Group-Aware<br>Constraint |
| --- | --- | --- | --- | --- |
| Full Framework | ✓ | ✓ | ✓ | ✓ |
| w/o Group-Aware | ✓ | ✓ | ✓ | - |
| End-to-End (CE) | ✓ | ✓ | - | - |
| Static Contrastive | - | ✓ | ✓ | ✓ |
| Single-Replica | ✓ | - | - | - |
| Static End-to-End | - | ✓ | - | - |

**Fig. 2.**
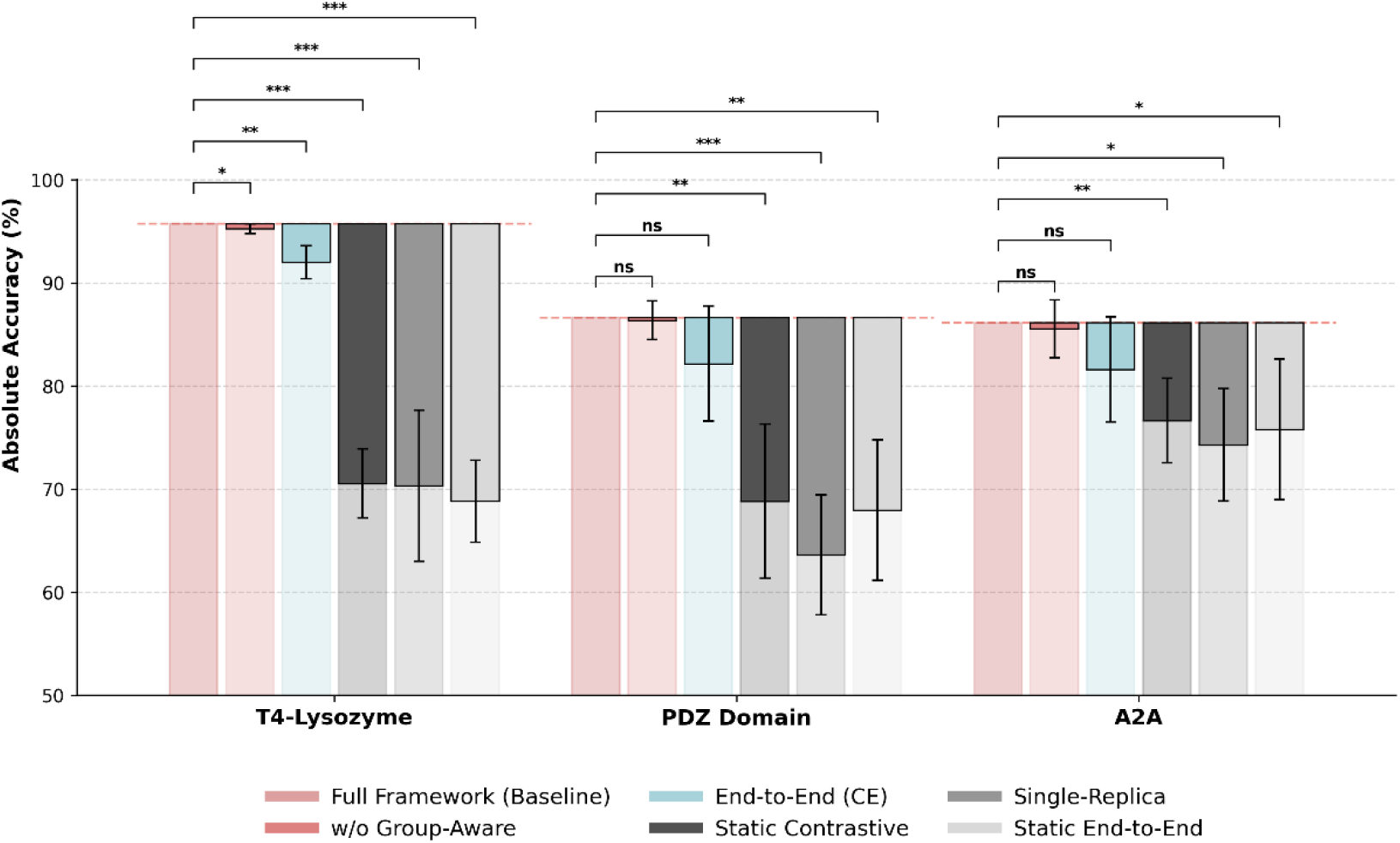
Ablation study demonstrating the cumulative impact of model components on classification accuracy. Performance of the model variants (components detailed in **Table 1**) was evaluated under a strict Leave-One-Replica-Out Cross-Validation (LORO-CV) regime to prevent trajectory-specific memorization. To explicitly visualize the functional loss caused by component ablation while accounting for the intrinsic biophysical variance of independent MD trajectories, a combined visual representation is employed. Semi-transparent bars indicate the absolute mean classification accuracy across independent held-out replicas (*n=5*). Solid colored bars highlight the exact performance gap, extending downward from the baseline reference (*Full Framework*, dashed red line) to the ablated variant’s accuracy. Accordingly, error bars denote the standard deviation of the paired, per-replica performance differences, strictly isolating the architectural impact from stochastic noise. Statistical significance was determined via one-sided paired *t*-tests against the baseline configuration (* *P* < 0.05, ** *P* < 0.01, *** *P* < 0.001, ns = not significant). The raw data are shown in the graph in **Extended Data Fig. 1.**

A fundamental bottleneck in applying deep learning to MD trajectories is the risk of overfitting to trajectory-specific patterns. Individual MD replicas are rarely fully ergodic and often retain the memory of their specific initial geometries, such as velocity distributions or localized structural drifts^25,27^. Deep neural networks, given their high representational capacity, can easily exploit these persistent patterns. We observed this limitation when training a baseline model in a Single-Replica setup. When evaluated on held-out frames from the same replica used for training, the model achieved an accuracy exceeding 99% (data not shown). However, when tasked to classify frames from a completely independent replica of the same target state, predictive performance declined sharply (**Fig. 2**, Single-Replica). The network had memorized the stochastic noise of the training simulation rather than learning the general dynamic signature of the state.

To overcome this trajectory dependence, GISTnet-MD employs a two-phase training protocol that decouples representation learning from the final classification task. While training the spatiotemporal model end-to-end with a standard cross-entropy loss (End-to-End CE) effectively filters a substantial amount of noise, our ablation analysis demonstrates that decoupling the training phases yields higher validation accuracies and greater robustness across independent starting conditions. In the first phase, the model uses contrastive learning to map temporal windows into a latent space according to their target state (**Fig. 1C**). To strictly prevent the model from exploiting intra-trajectory correlations, we introduced a group-aware triplet selection strategy. Instead of pairing arbitrary windows from the same state, the algorithm aligns an “anchor” window with a “positive” window sampled strictly from a different, independent MD replica. Simultaneously, it maximizes the distance from a “negative” window sampled from the opposing target state. This mapping explicitly penalizes reliance on replica-specific noise, driving the model toward the learning of state-associated biophysical features. The ablation study confirms the impact of this design (**Fig. 2**). While removing the explicit group-aware sampling constraint (w/o Group-Aware) yielded only minor changes in performance, the constraint was retained in the Full Framework to algorithmically enforce the neutralization of initial-condition bias across independent replicas. Once the latent space is optimized, the spatiotemporal backbone is frozen. In the final phase, a linear classification head is trained on these fixed embeddings to obtain the final state predictor. Furthermore, applying contrastive learning without temporal integration (**Fig. 2**, Static Contrastive) leads to a drop in accuracy. This confirms that the contrastive loss alone is insufficient, and the explicit modelling of temporal data is crucial to build a functional latent space.

Finally, to translate the optimized latent representations from the model into biophysical insights, the architecture incorporates an explainable AI (xAI) pipeline. Specifically, an IG algorithm is applied to compute the gradients of the model’s prediction confidence with respect to the input spatial features. This approach effectively maps the network’s latent decision boundaries back to specific three-dimensional residues and interactions.

### Estimating Thermodynamic Sampling Convergence

By systematically evaluating the number of independent trajectories required for GISTnet-MD to generalize to unseen data, it is possible to assess when the model has successfully extracted the invariant patterns defining the target state. To this end, the number of independent MD replicas used during the training phase was progressively increased from 1 to 8 and performance was monitored for the T4-Lysozyme and PDZ domain systems (**Fig. 3**).

**Fig. 3.**
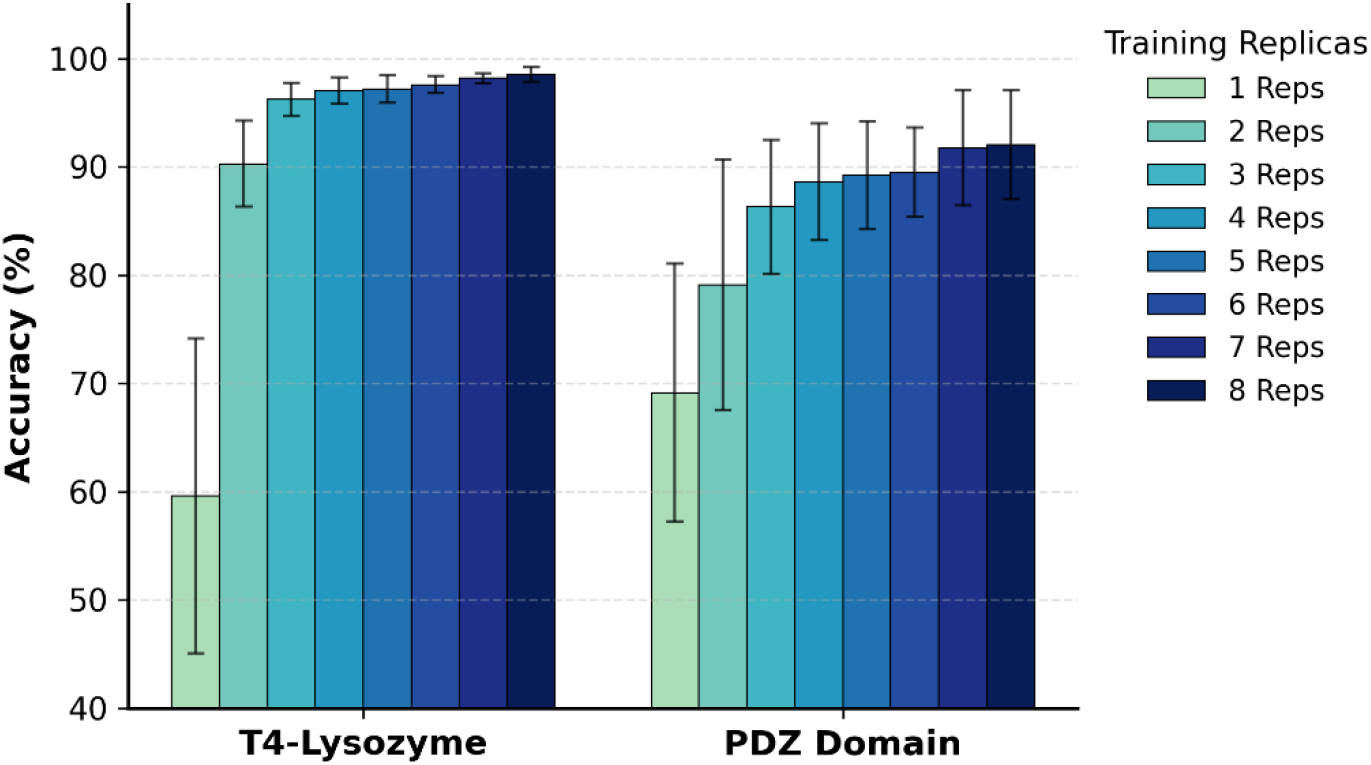
Quantifying thermodynamic sampling convergence via data scaling. Evaluation of model performance as a function of training data volume across the T4-Lysozyme and PDZ Domain systems. Bar charts display the mean validation accuracy achieved when training on an increasing number of independent MD replicas (from 1 to 8 replicas). Bars represent the mean accuracy calculated across independent validation folds (n=5). Error bars denote the raw standard deviation across these folds\The sequential color gradient indicates the progressive inclusion of additional independent MD replicas in the training set..

The A2A receptor case was excluded from this analysis due to the limited number of independent agonist-bound simulations available in the GPCRmd database^41^. Evaluations were performed using a 5-fold cross-validation scheme (raw data is shown in the graph in **Extended Data Fig. 2**).

The performance at increasing number of trajectories demonstrates a distinct convergence behaviour. When the training set is restricted to a minimal number of replicas (e.g., 1 or 2), average accuracy remains low and is accompanied by high variance, consistent with the model overfitting to stochastic thermal noise and with limited sampling diversity. As the number of training replicas increases, validation accuracy improves, reaching a stable plateau. This plateau indicates that the model has reached information saturation; beyond this threshold, the introduction of additional independent trajectories does not provide novel, state-specific information. The exact number of replicas required to reach this saturation is system dependent. For localized perturbations, such as the cavity-forming mutation in T4-Lysozyme, information saturation is achieved rapidly, exhibiting a plateau around 3 to 4 replicas. For the PDZ domain, which is characterized by distributed allosteric dynamics and higher inter-replica variance, a larger number of independent trajectories is required. In both scenarios, convergence implies that a finite set of independent replicas is sufficient to capture the biophysical signature of the conformational ensembles. Consequently, this data scaling approach provides a data-driven metric to estimate the sampling depth required for GISTnet-MD to adequately capture the functional differences between two macroscopic states.

### Explainable AI: Localized gating in T4-Lysozyme

To evaluate the mechanistic interpretability of GISTnet-MD, the framework was applied to bacteriophage T4-Lysozyme (T4L), comparing the wild-type (WT) and L99A mutant^29–31^ dynamics. The L99A substitution generates a buried hydrophobic cavity of approximately 150 Å^3^, shifting the conformational ensemble toward a binding-ready state. Geometric analysis of the simulation trajectories confirmed the presence of a localized cavity (**Supplementary Note 2**). However, standard global geometric metrics (including RMSD, RMSF, and principal component analysis) did not clearly separate the WT and mutant ensembles (**Extended Data Fig. 3**). Both states populate overlapping conformational basins and exhibit comparable global fluctuation profiles, with only minor RMSF deviations observed between residues 70 and 90. This prevents the isolation of the functional mutant signature through standard analytical methods.

To assess if GISTnet-MD can overcome this limitation without relying on the label of the mutated site, residue 99 was explicitly excluded from the input graph (i.e., its corresponding node and spatial edges were completely removed). The network was consequently forced to identify the target state solely through the dynamic propagation of the perturbation to the surrounding residues. Structural mapping of the consensus Integrated Gradients (IG) scores indicates that the model focuses on residues flanking the mutation site (**Fig. 4**). The highest attribution score is assigned to I78, a residue located in the C-terminal domain helix that contributes to the hydrophobic core and lines the cavity, alongside L84, M102, L118, L121, and L133.

**Fig. 4.**
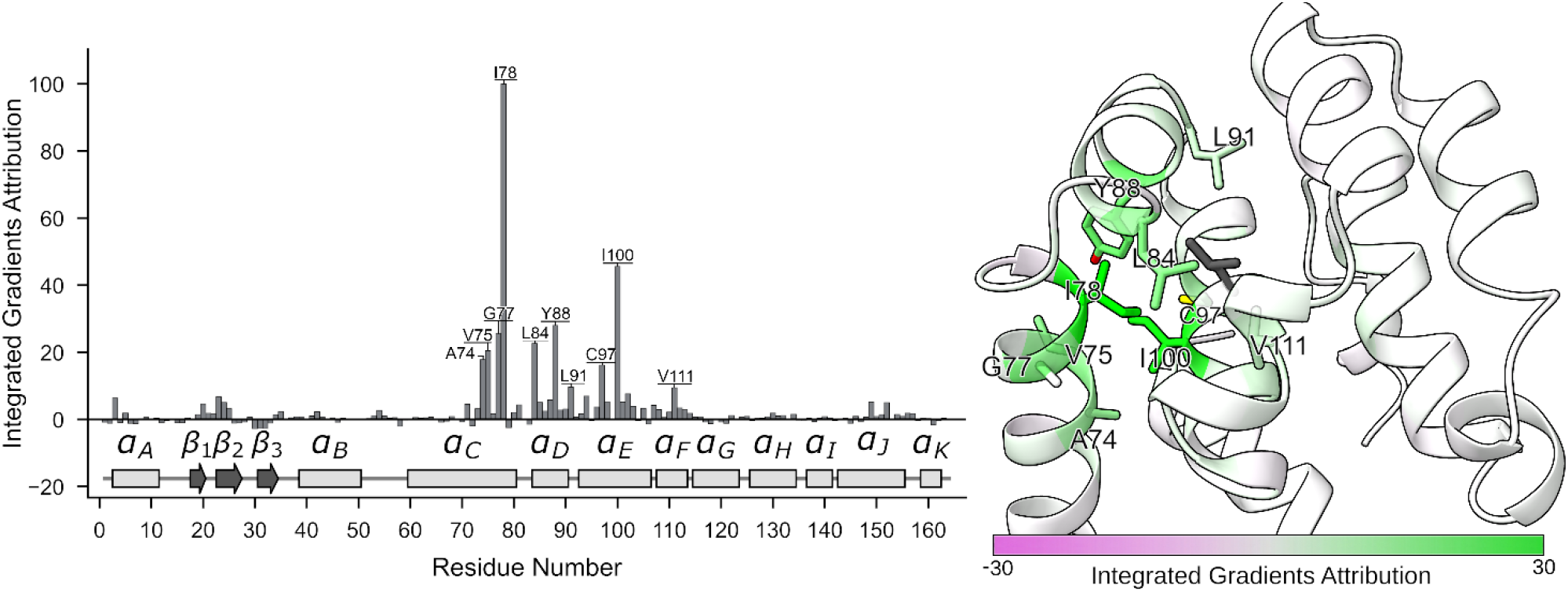
Explainable AI identifies the gating mechanism of the T4-Lysozyme binding cavity. **a**, Residue-level attribution plot. The bar height represents the consensus IG score for each residue. The protein’s secondary structure elements are mapped along the x-axis for topological reference. Labels indicate the 10 residues with highest IG attribution. **b**, 3D structural mapping of the consensus Integrated Gradients (IG) scores on the T4-Lysozyme L99A mutant (PDB ID: 3DMV). Residues are colored according to their aggregated importance. The mutated residue 99 is represented as gray sticks for reference but was excluded from the input graph. The top 10 residues contributing to the classification are shown in stick representation and labeled.

Mechanistically, I78 participates in the gating of the cavity. Prior MD studies indicate that ligand entry requires transient openings between helices Dα and Gα^42^. As previously described, in the L99A mutant, the reduced steric bulk of A99 permits the I78 side chain to shift toward position 99, lowering the energetic barrier for the helical rearrangements required for access. In the WT protein, the L99 sidechain induces a steric clash with I78 during these motions, penalizing the open conformations and reducing their probability. By identifying I78 as the primary perturbed node, GISTnet-MD successfully recovers this known gating mechanism directly from the raw trajectory data.

While experimental literature documents the role of the Fα and Gα helices in T4L dynamics, specifically concerning a conformational switch of F114^43,44^, the model assigns lower importance scores to this region. Only V111 ranks among the top 10 residues for IG attributions. This occurs because the F114 flip was not sampled within the simulation trajectories. In the absence of this transition, the network bases its classification on the dynamic signatures of cavity breathing and gating localized at the cavity interface adjacent to the masked mutation site.

### Explainable AI: Distributed allostery in the PDZ3 domain

The application of the framework to the PDZ3 domain evaluates the ability of GISTnet-MD to recognize dynamic states associated with peptide binding. This domain represents a classic model of dynamic allostery, where ligand binding modulates distal sites altering the protein’s conformational fluctuations, without inducing macroscopic structural rearrangements. Standard molecular dynamics metrics indicate the C-terminal region as the most flexible region of the domain (**Extended Data Fig. 4**) without significative difference between apo and holo states. IG scores from the model trained only on the protein graph (excluding the peptide) show that the determinants for discriminating between the apo and holo states are the residues of the peptide-binding site, specifically the αB helix and the β2 strand (**Fig. 5a,d**). The β3 strand, which is not a primary contact site, exhibits elevated attribution scores. Interestingly, the network tracks partial unbinding in time-resolved detail, updating its state predictions and IG attributions to reflect the instantaneous loss of intermolecular contacts (**Supplementary Note 3**). To assess whether the network identifies distributed allosteric coupling rather than local dynamic alterations due to the steric hindrance of the bound peptide, an *in silico* ablation experiment was conducted. A second model was trained on the same simulations but with the binding site masked (removing all residues belonging to the αB helix, the β2 strand, and their contiguous regions from the input graph). This approach allows us to silence the primary signal and force the network to focus on the more subtle, distributed dynamic couplings. The masked model averaged 60% accuracy, with cross-validation fluctuations often approaching 50% (**Fig. 5b**). Expanding the training dataset to ten MD replicas (eight for training, two for validation) increased the masked model’s average accuracy to 76%, maintaining a baseline above 60% across all cross-validation folds (**Fig. 5b**). The increase in performance at the increase of training data suggests that in systems exhibiting elevated structural fluctuations and substantial inter-replica variance, the subtle dynamic signature may be partially obscured. The integration of additional independent replicas increases statistical sampling improving the signal/noise ratio for the allosteric signature. The successful classification of the target state in the absence of the primary interaction site show that the dynamic signature can be detected independently from information on local ligand-induced changes.

**Fig. 5.**
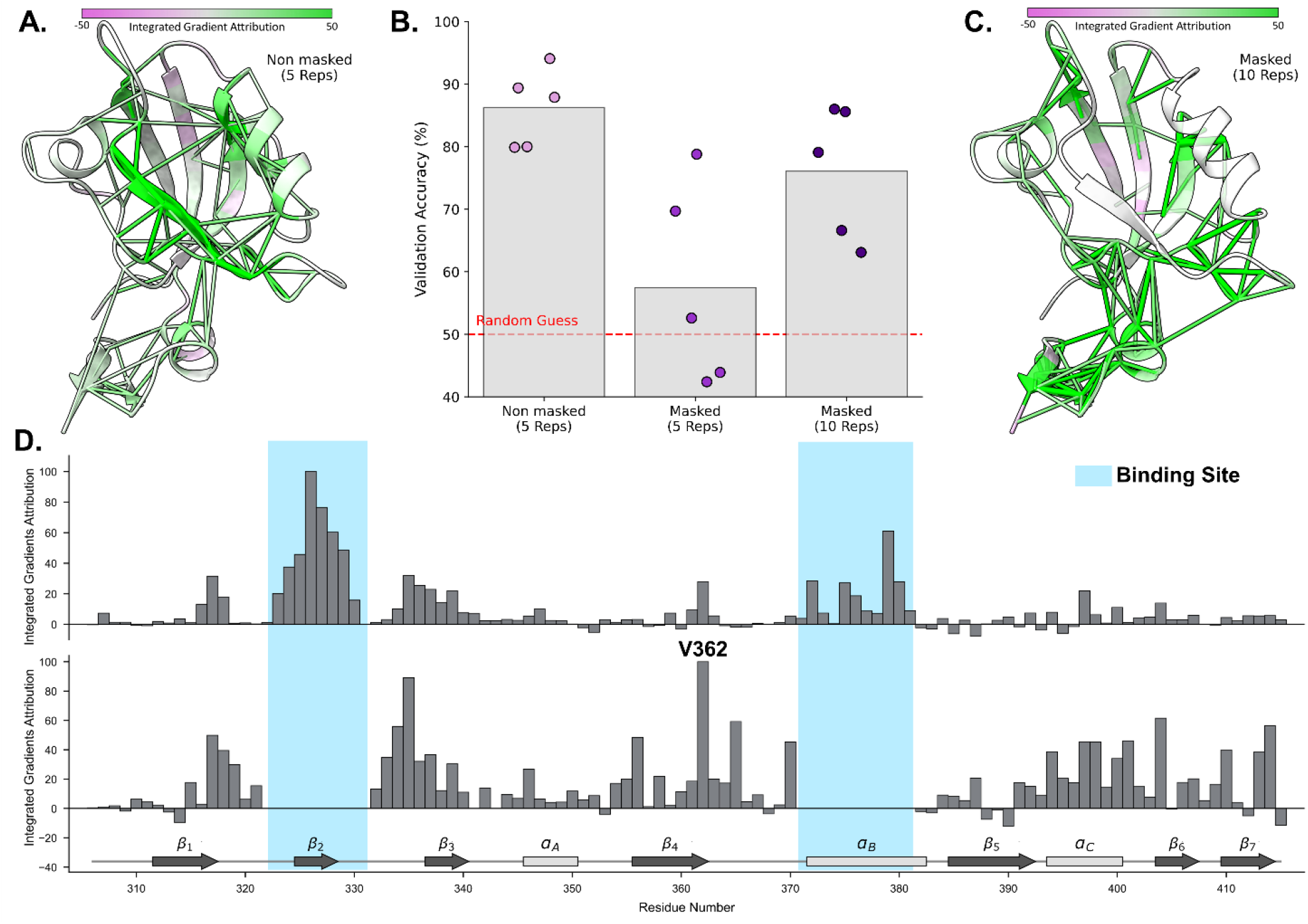
Isolating distributed dynamic allostery in the PDZ3 domain via structural masking. **A:** Network visualization of the dynamic interactions determining the apo versus holo classification in the standard (non-masked) GISTnet-MD model. Residues are colored according to the per-residue IG attribution score. The 100 highest-scoring edges for classification are represented as cylinders, with thickness and color intensity scaled to the edge-specific IG scores. **B:** Impact of structural masking and sampling on model convergence. The bar plot reports the classification accuracy for the complete model (Non-masked, 5 Reps), the model trained without the binding site residues (Masked, 5 Reps), and the model trained without the binding site residues utilizing an expanded dataset of 10 independent MD replicas (Masked, 10 Reps). Bars denote the mean validation accuracy; dots represent the accuracy of independent cross-validation folds. The dashed line indicates the 50% threshold. **C** 3D IG scores of the masked model. Residues and the top 100 high-scoring edges are colored according to their IG attribution scores, derived from the Masked (10 Reps) model. **D:** Spatial distribution of the thermodynamic attribution. Stacked 1D bar charts display the per-residue consensus IG scores for the non-masked (top) and masked (bottom) models. Secondary structure elements are indicated along the x-axis. Shaded areas denote the primary peptide-binding regions (β2 strand and αB helix) omitted from the input graph in the masked configuration.

The IG attribution analysis of the masked model can be used to map the residue network associated with the allosteric communication (**Fig. 5c, d**). Upon masking the primary binding site, the highest attribution scores shift to the β3 strand and the C-terminal region. However, a number of residues across the domain also display local dynamic alteration, suggesting that the allosteric signal is diffuse. Specifically, the absolute maximum IG attribution peak is located at Val362, situated on the distal face of the β-sheet (β4). These computational results are consistent with two distinct experimental results documented in the literature. First, thermodynamic and NMR analyses demonstrated that peptide binding modifies the structural dynamics of the C-terminal region, encompassing the αC elements^32,33,35,45^. Second, independent perturbation response scanning and NMR studies indicated that multiple distributed domain residues, explicitly including Val362, act as discrete sites undergoing local dynamic alterations upon ligand recognition^32,33,46^. This agreement supports GISTNet-MD’s potential to isolate the subtle allosteric signatures that define a functional state.

### Explainable AI: Activation signatures in the Adenosine A2A GPCR

To evaluate GISTnet-MD ability to detect general activation signatures across chemically diverse ligands, the framework was applied to the Adenosine A2A GPCR. Simulation trajectories were retrieved from the GPCRmd database, including conformations bound to four distinct agonists (active state) and four distinct antagonists (inactive state). The initial comparative task classified the agonist-bound versus antagonist-bound MD ensembles. To ensure the extraction of general activation signatures independent of specific ligand chemistries, model evaluation was executed using a Leave-One-Group-Out Cross-Validation (LOGO-CV) scheme. In this protocol, an entire ligand group (including all associated independent replicas) was excluded during training and used exclusively for validation. Because these simulations originate from distinct experimental crystal structures (active versus inactive), they contain inherent geometric disparities, including the macroscopic conformational shift of the intracellular half of Transmembrane Helix 7 (TM7) and the associated NPxxY microswitch (**Extended Data Fig. 5a**). Consistent with this structural difference, the unmasked model’s IG scores are consistent with this structural divergence (**Extended Data Fig. 5b, c**). To assess if the model captured distributed allosteric coupling rather than the difference in conformational endpoints, a structural masking protocol was performed. A masked model was trained, restricting the input graph to residues 1-280, thereby removing the intracellular segment of TM7 and the NPxxY motif (**Extended Data Fig. 6a**). The masked network retained excellent classification performance (**Fig. 6a**). Following the removal of the primary macro-switch, the IG scores highlighted an intracellular hydrophobic cluster bridging TM5 and TM6. The network assigned high IG scores to residues V196, Y197, I200, and F201 on TM5, and L235, I238, and V239 on TM6 (**Fig. 6b, d**). This region corresponds to the hydrophobic hinge coordinating the outward swing of TM6, an established hallmark of Class A GPCR activation^37,39,47,48^. The identification of this secondary allosteric hub, where inter-state structural divergence is minimal and comparable to the intrinsic variance among the starting agonist-bound structures, indicates the extraction of activation-associated dynamic signatures that extend beyond the dominant macroscopic structural divergence.

**Fig. 6.**
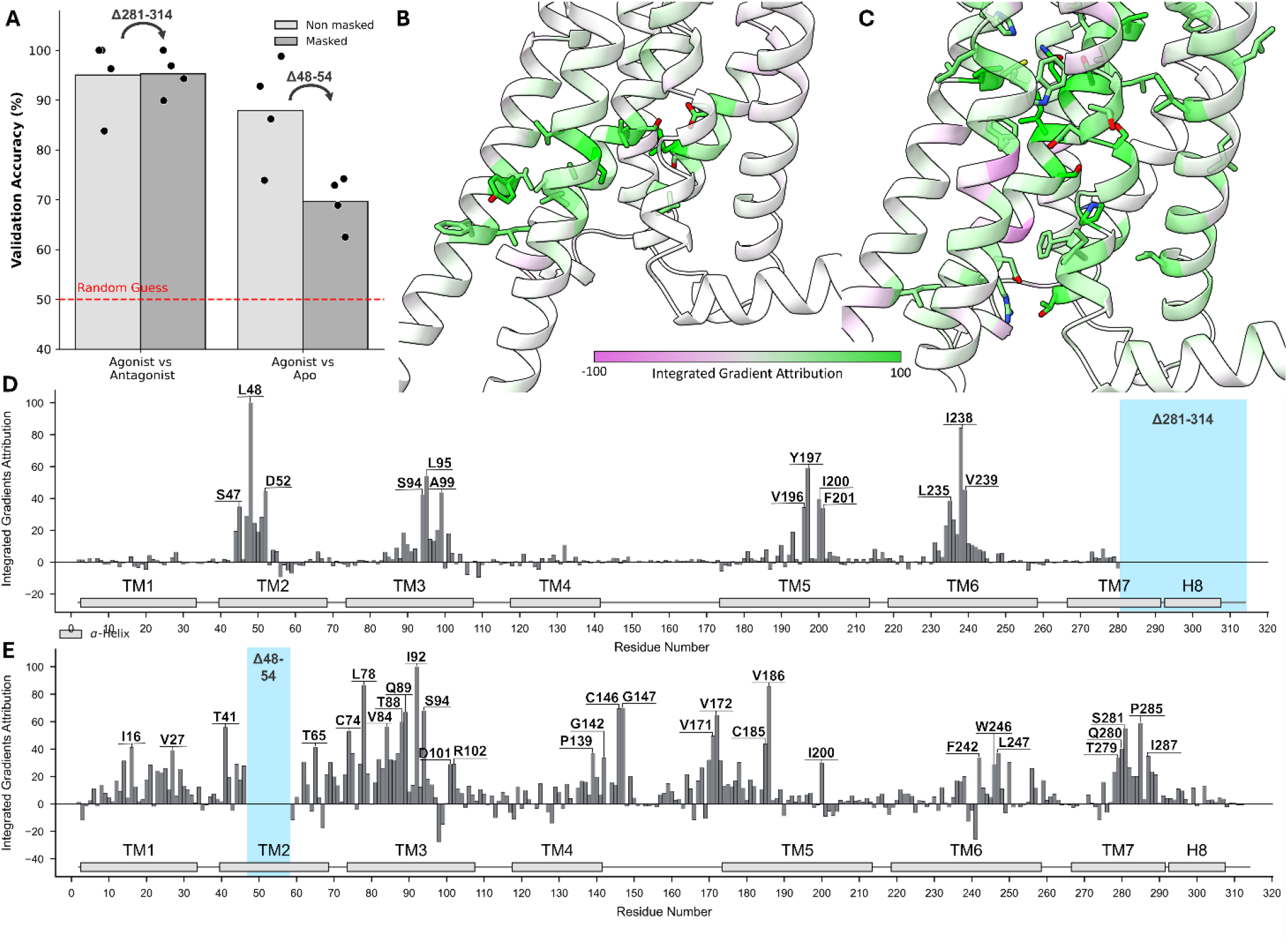
Decoupling structural biases to isolate allosteric activation signatures in the Adenosine A2A GPCR. **A:** Impact of structural masking on classification accuracy for two comparative tasks: Agonist vs. Antagonist (differentiating ensembles originating from active and inactive crystal structures) and Agonist vs. Apo (evaluating differential relaxation dynamics from identical active coordinates). Bar plots report the mean validation accuracy for the complete (non-masked) and masked models across independent cross-validation folds (dots). The dashed line indicates the 50% threshold. **B:** Structural attribution of the Agonist vs. Antagonist classification after masking the TM7 intracellular segment (residues >280), mapped onto the A2A 3D structure. **C:** Structural attribution of the Agonist-bound vs. Apo classification after masking the electrostatic bias of the sodium pocket (residues 48–54). **D:** 1D consensus IG profile for the Agonist vs. Antagonist classification following the masking of the TM7 intracellular segment. **E:** 1D consensus IG profile for the Agonist-bound vs. Apo classification following the masking of the sodium pocket. Transmembrane helices are indicated along the x-axis in **D** and **E**.

In addition, to isolate the dynamic signature induced by agonists binding, we compared agonist-bound (active-holo) trajectories against active-apo trajectories (MD simulations started from the same agonist-bound conformation but with the ligand removed). This setup limits the classification task to detecting the signature of relaxation dynamics prior to the global structural deactivation. Initial analysis indicated that the network identified a specific difference in the MD simulation set-up: the differential protonation state of D52 in the sodium pocket (**Extended Data Fig. 7**). While this electrostatic shift is a well-documented hallmark of the active state^36^ and was deliberately incorporated into the simulation protocols of the GPCRmd database^41^, its dominant signal obscures the underlying activation signal. To decouple this allosteric signal from the D52 electrostatic perturbation, an additional structural masking step was performed by excluding the sodium pocket (residues 48–54, **Extended Data Fig. 6b**). The masked model displayed a drop in accuracy but retained a discrete cross-validation accuracy of approximately 70% (**Fig. 6a**). Following the masking of D52, the IG attribution shifted to the lower region of the orthosteric pocket, reflecting the loss of ligand interactions (**Fig. 6c,e**). High IG scores were assigned to W246, the toggle switch probing the agonist’s ribose moiety^37,47,49^, and F242, the core of the PIF motif^36,47^, along with their surrounding residues such as Q89, I92 and V186. The attribution network traces the established allosteric signal propagation along TM3, with peaks on I92 (of the PIF motif^36^), Q89 and T88 connecting the orthosteric sensors to the intracellular space^37,50^. The attribution extends to the intracellular effectors, yielding signals on P285 of the NPxxY motif (TM7), D101 and R102 of the DRY motif (TM3), and T41 at the G-protein coupling interface^36,37,39,47^.

Rather than relying on the full global structural deactivation, which may require timescales beyond standard simulation limits, GISTnet-MD was tested on 500-ns trajectories but successfully captured the immediate relaxation dynamics following ligand removal. By processing the simulations across multiple independent replicas, the network filters out stochastic transient fluctuations and is able to detect the initial dynamic uncoupling. This leads to the identification of early deactivation-associated dynamics, successfully tracking the initial allosteric signal propagation from the orthosteric switches to the intracellular microswitches. It should be noted that this is based on a dynamic signature, not requiring information on the large-scale structural changes.

## Discussion

Extracting the functional dynamical signatures that define a binding state from the stochastic noise of molecular dynamics simulations still represents a fundamental challenge in computational biophysics. Traditional analysis methods, relying on geometric variance or linear dimensionality reduction, frequently fail to isolate functional differences when the dynamic signature is subtle, delocalised, or allosteric. GISTnet-MD overcomes these limitations with a framework that integrates continuous graph representations, spatiotemporal aggregation, and group-aware contrastive learning. The combination of these methods algorithmically encourages the model to isolate replica-invariant biophysical patterns, effectively decoupling generalized thermodynamic differences from initial coordinate biases and trajectory-specific noise.

The robustness and generality of this methodology is demonstrated with examples of dynamic perturbations of varying complexity. To remove initial biases, molecular systems were initialized from identical coordinates. In all cases, GISTnet-MD successfully extracted subtle difference in relaxation dynamics across all systems. In the localized perturbation of T4-Lysozyme, where the conformational ensembles have overlapping global fluctuations, the network isolated the signature motion of the cavity gating mechanism (I78 steric clash) even if the mutation site was explicitly masked. In the PDZ3 domain, the trained model revealed long-range dynamic allostery, tracing communication pathways to the distal β3 strand and the C-terminal αC helix, without information on the primary steric hindrance of the ligand. In the Adenosine A2A GPCR, the model identified functional signatures across comparative tasks of varying complexity and initial conditions. When comparing active and inactive ensembles originating from divergent structures, the model captured both primary dynamics differences and, following their *in silico* ablation, the underlying generalized allosteric propagations. Additionally, when comparing agonist-bound and active-apo trajectories to classify differential relaxation dynamics, the model successfully isolated the distributed allosteric network associated with early receptor deactivation. In all these examples, we also demonstrated that a rigorously baselined Explainable AI (xAI) can return interpretable information on the structural determinants of functional dynamics at the residue and interaction level. These results can then be directly used to plan mutational scanning and pharmacological targeting.

The ability to extract structural hypotheses is rooted in the spatiotemporal and invariant design of GISTnet-MD. Recent data-driven approaches, such as DiffNets^16^, have successfully used self-supervised autoencoders to map structural ensembles to biochemical properties by processing static arrays of Cartesian coordinates. GISTnet-MD extends this paradigm to dynamic representations. By encoding spatial topology through continuous graphs and integrating dynamics via 1D-CNN sliding windows, the model architecture preserves biophysical patterns and filters high-frequency noise. The choice of robust deep learning components is instrumental in ensuring statistical stability when processing noisy MD datasets, preventing the model from overfitting to stochastic fluctuations. Similarly, in allosteric analysis, recent GNN architectures like AlloPool^21^ have introduced an autoencoder paradigm that learns to reconstruct the dynamics, using temporal attention to iteratively prune residue interactions and directly identify minimal allosteric networks. Unlike models optimized for unsupervised reconstruction, GISTnet-MD relies on predicting target state labels. By employing a group-aware contrastive pre-training phase, our model decouples representation learning from the classification task. This guides the optimization toward generalized biophysical invariants removing the initial condition biases and replica-specific noise. Furthermore, rather than extracting biological insights through structural pruning, GISTnet-MD preserves the complete protein graph topology, and the mechanistic interpretation is delegated to the final Explainable AI (xAI) pipeline based on an Integrated Gradients (IG) algorithm.

However, the successful extraction of functional determinants is dependent on the underlying simulation dataset being representative of the process under study. Historically, determining whether a simulation has adequately sampled a functional state relies on heuristic geometric criteria. By mapping accuracy against the number of independent replicas, GISTnet-MD offers an alternative strategy to evaluate the convergence of state-discriminative information in a set of trajectories. Data scaling analysis suggests that the convergence threshold is dependent on the specific biophysical feature under evaluation. For example, while five independent replicas provided sufficient statistical power to classify the PDZ3 domain based on the primary peptide-binding interface, this proved inadequate when using *in silico* masking of the binding site. In this ablated regime, where the network was constrained to extract a diffuse, allosteric signal from high inter-replica structural variance, generalization of the accuracy degraded. Robust classification was only achieved by expanding the dataset to ten independent replicas. In this context, the emergence of an accuracy plateau indicates a state of predictive convergence. While this does not guarantee that the free-energy landscape has been exhaustively explored, it confirms that the simulation has captured sufficient variance to robustly isolate the distinguishing functional signature. Consequently, GISTnet-MD offers an objective, data-driven metric to evaluate simulation convergence based directly on the extracted functional signature, rather than relying on surrogate geometric variables. More generally, this data scaling test confirmed a well-known practical principle for molecular dynamics: increasing the number of independent replicas is often more informative than extending a single trajectory. While long simulations are sometimes needed to cross energy barriers, capturing the consistent fluctuations that define conformational ensembles requires diverse initial velocities to prevent the analysis from overfitting to replica-specific noise.

Despite its potential, GISTnet-MD exhibits methodological limitations worthy discussing. First, it is strictly data-dependent; the network cannot discover rare conformational events (such as the Phe114 flip in T4-Lysozyme) if they are not explicitly sampled within the input trajectories. It functions as an analyser of existing structural variance rather than an enhanced sampling technique or a generative framework. Second, timescale misalignment remains a constraint. In the A2A active-apo simulations, 500 ns may be insufficient for complete macroscopic deactivation, so the model detects the initial dynamic uncoupling triggered by ligand removal but cannot extrapolate the full thermodynamic pathway unless the simulation physically traverses it. Third, the extraction of stable biophysical functional patterns depends upon the availability of a substantial volume of independent trajectories. As evidenced by the PDZ3 masking experiments, resolving subtle allosteric signals requires an increased number of independent replicas to effectively suppress stochastic fluctuations. Fourth, investigating complex allosteric or distributed signal currently relies on an iterative masking procedure. While sequentially masking dominant perturbation sources (such as the primary binding interface in PDZ3 or the TM7 shift in A2A) isolates localized thermodynamic signals, this iterative ablation increases computational cost. To overcome this limitation, future developments will focus on diversifying the mechanism of feature extraction to resolve these secondary components of the functional network without the need for explicit masking. Finally, the paradigm is inherently comparative, requiring the explicit sampling of at least two distinct states to define the contrastive objective.

In conclusion, GISTnet-MD establishes a physically grounded framework for the comparative analysis of molecular dynamics trajectories sampled across distinct conformational ensembles. Isolating invariant dynamic signatures from stochastic noise, this spatiotemporal deep learning framework provides a methodology to decode the structural and dynamic features underlying biomolecular regulation. In principle, GISTnet-MD can be generalized to a broader range of comparative MD applications, opening new avenues for evaluating the functional impact of post-translational modifications, varied protonation states, or different environmental conditions.

## Methods

### The GISTnet-MD architecture

#### Operational Framework and Label Definition

The extraction of invariant dynamic signatures is formulated as a discrete classification task operating on temporal windows of MD trajectories. In our comparative framework, independent MD simulations are explicitly generated to sample the conformational ensembles of distinct macroscopic biological conditions (e.g., apo versus holo, or WT versus mutant). To provide the objective targets for the model’s training phase, we employ a trajectory-wide labeling strategy. We operate under the assumption that the independent replicas consistently sample the biological state they were initialized to represent. Consequently, every sliding temporal window extracted from a given simulation replica strictly inherits the macroscopic label of its parent condition. For instance, all temporal windows extracted from an “apo” simulation are uniformly assigned the target label “apo”, while those from a “holo” trajectory are labeled “holo”. By tasking the neural network to predict these predefined discrete labels from the continuous trajectory data, the model is forced to identify the underlying structural and dynamic features that consistently differentiate the chosen target ensembles.

#### Continuous Graph Representation

To capture both residue interactions and time-dependent structural changes, Molecular Dynamics (MD) trajectories are discretized into a sequence of coarse-grained residue-residue graphs. For a given frame *t*, protein coordinates are converted to an undirected graph, where each node represents an amino acid residue and each edge a pair of interacting residues. To prevent label leakage, graphs were constructed utilizing protein residues only; ligand molecules, ions, or bound peptides were explicitly excluded from the graph nodes and features. Furthermore, in mutational studies (e.g., T4-Lysozyme L99A), the mutated residue identity was fully masked from the input graph. Nodes are initialized with a one-hot encoded vector 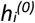 representing the residue identity. Edges are drawn between any node pair (*i, j*) whose Euclidean distance d_ij_ falls within a predefined interaction cutoff *r*_*cut*_ (e.g., 10 Å between their Centre of Mass in this study). To provide a richer spatial representation, scalar distances are expanded using a set of Gaussian Radial Basis Functions (RBFs):

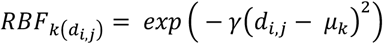

where *µ*_*k*_ are centers linearly spaced between 0 and *r*_*cut*_. Moreover, these RBF features are modulated via multiplication with a continuous envelope function *U(d*_*ij*_) to prevent sudden discontinuities when residue-pair distances approach or cross the cutoff boundary. Following the formulation introduced in DimeNet^51^ we employed a polynomial envelope that ensures both the function and its spatial derivatives smoothly decay to zero at the cutoff boundary. Defining the normalized distance *x = d*_*ij*_*/r*_*cut*_, the envelope is:

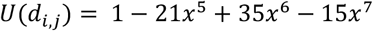

for *d*_*ij*_ *< r*_*cut*_, and *U(d*_*ij*_*) = 0* otherwise. The final edge feature vector is therefore given by the element-wise product of the RBF expansion and the cutoff envelope:

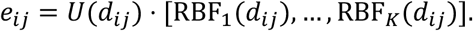

This representation provides smooth distance-dependent edge features for the continuous-filter graph neural network.

#### Spatial Encoder (Continuous Filter Convolution)

Structural embeddings are then extracted from single frames using a message-passing neural network inspired by SchNet^40^. This encoder applies continuous-filter convolutions in which the interaction weights are dynamically generated based on the RBF-expanded inter-residue distance features e_ij_. In this way, messages between neighbouring residues are modulated by their spatial separation, allowing the model to learn local structural environments and distance-dependent residue interactions. The output of the spatial encoder is a learned residue-level embedding for each frame, which is subsequently passed to the temporal module described below.

#### Spatiotemporal Integration (1D-CNN)

*The embedding tensor is reshaped so that each residue trajectory is treated as an independent temporal sequence with D input channels. The 1D-CNN therefore learns temporal filters over the sequence of GNN-derived residue embeddings*.

*The temporal module consists of two stacked convolutional layers with kernel size k=3. The first layer preserves temporal resolution using stride s=1, while the second layer performs temporal downsampling using stride s=2. Each convolutional layer is followed by batch normalisation, SiLU activation, and dropout regularisation. A final adaptive average pooling layer compresses the remaining temporal dimension, producing a single time-aware embedding h_i^’ ∈R^ D for each residue i within the window*.

To capture local fluctuations *while reducing sensitivity to short-timescale* thermal noise, GISTnet-MD processes sliding temporal windows of length *T* (Fig 1B). For each window, the spatial encoder first converts each frame independently into residue-level embeddings, producing a tensor of shape *B, T, N, D*, where B is the batch size, N the number of residues, and D the hidden dimension (*D=128* in our baseline configuration). Temporal integration is then performed using a 1D-CNN. The temporal convolution is applied independently to the temporal history of each residue and does not pool across the residue graph.. The tensor is reshaped so that each residue trajectory is treated as an independent temporal sequence with *D* input channels. The 1D-CNN therefore learns temporal filters over the sequence of GNN-derived residue embeddings. The temporal module consists of two stacked convolutional layers with kernel size k=3. The first layer preserves temporal resolution using stride s=1, while the second layer performs temporal downsampling using stride s=2 (**Fig. 1B**). Each convolutional layer is followed by batch normalization and SiLU activation. A final global average pooling layer compresses the remaining temporal dimension, producing a single time-aware embedding *h’*_*i*_ (of dimension *D*) for each residue *i* within the window.

*Global Attention Pooling:* To compress the residue-level dynamic fingerprints into a single state vector representing the entire temporal window, a Masked Global Attention Pooling mechanism is used. An additive attention network (a two-layer MLP with Tanh activation) computes an unnormalized importance score *s*_*i*_ for each residue embedding *h’*_*i*_:

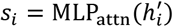

The attention weights are then obtained by applying a softmax over the residue dimension:

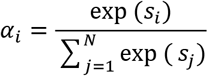

where masked or excluded residues are omitted from the normalisation. The final graph-level embedding *Z* for the temporal window is computed as the attention-weighted sum of the residue embeddings:

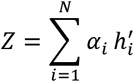

This pooling step allows the model to assign greater weight to residues whose temporal embeddings contribute most strongly to the state representation, while preserving a fixed-dimensional representation for downstream contrastive learning and classification.

#### Group-Aware Contrastive Learning and Linear Probing

Training end-to-end classification models on highly correlated MD trajectories can lead to overfitting, as the network tends to memorize replica-specific correlations rather than learning the biophysical features that generalise across independent simulations. To reduce this risk, GISTnet-MD is trained using a two-phase protocol.

##### Phase 1: Representation Learning

The model is first trained to organise temporal-window embeddings according to state-level similarity using a *Group-Aware Contrastive Loss* (**Fig. 1C**). Given a batch of window embeddings, an Anchor (*A*) and a Positive (*P*) are defined as a pair of windows belonging to the *same* functional state (e.g., Apo) but strictly sampled from *different* independent MD replicas (groups). A Negative (*N*) is defined as a window sampled from a different functional state. The loss function reduces the Euclidean distance between *A* and *P* while separating *A* from *N* by at least a predefined margin *m*. This forces the model to map frame windows from independent replicas of the same state to occupy the neighbouring regions of the latent space, while reducing reliance on replica-specific correlations.

##### Phase 2: Linear Probing

After contrastive pre-training, the weights of the spatiotemporal backbone are frozen. A simple Linear Classifier (a BatchNorm layer followed by a single Linear layer) is then trained on the fixed embeddings using standard Cross-Entropy Loss to define the final decision boundary between target state labels.

Model’s architectural and temporal parameters were selected after a comprehensive hyperparameter analysis to obtain an optimal balance between predictive accuracy and computational tractability (see **Supplementary Note 4**).

### Leave-One-Group-Out (LOGO) Cross-Validation

To assess wether GISTnet-MD generalises to unseen trajectories, model evaluation, hyperparameter tuning, and early stopping were performed using a Leave-One-Group-Out Cross-Validation (LOGO-CV) regime. The dataset was partitioned into discrete groups where the definition of a group depended on the biological comparison being evaluated. For each fold, the model was trained on a subset of these groups and validated on a completely held-out group that is strictly excluded from training. Training and validation are repeated iteratively, each time holding out a different group, until each group had served once as the validation set. For systems where the target state is sampled via multiple independent simulation runs (e.g., T4-Lysozyme and the PDZ3 domain), a group is defined as a single MD replica. In this scenario, the LOGO-CV protocol computationally equates to a Leave-One-Replica-Out (LORO-CV) scheme. For systems in which multiple chemically distinct perturbations represented the same macroscopic state (e.g., the Adenosine A2A GPCR activated by diverse agonists), each group contained all independent simulation replicas associated with a specific ligand. This configuration corresponds to a Leave-One-Ligand-Out protocol and evaluates whether the learned representation transfers to ligand chemistries not observed during training.

### Explainable AI (xAI) and Structural Attribution

To interpret the physical determinants driving the classification, an xAI pipeline based on the IG method was implemented^52^. This approach maps the learned decision boundaries back to specific temporal frames and topological residue-level interactions. For any given temporal window, the model’s prediction confidence is defined as the softmax probability output of the linear probe classifier. The IG algorithm computes the gradients of the model’s output with respect to the input edge features (i.e., the RBF-expanded inter-residue distances) relative to a baseline.^52^ To provide a rigorous biophysical contrast, we adopted an Expected Gradients approach, explaining the prediction of a target class by contrasting it against a reference thermodynamic ensemble from the opposing classes. This procedure is articulated in the following steps:

1. *Reference Pool Selection:* Windows correctly classified with a strict confidence threshold (> 80%) are filtered to ensure the purity of the reference ensemble.
2. *Latent Clustering for Baseline Selection:* Computing Expected Gradients across all frames of the opposing class is computationally prohibitive. To obtain a limited yet comprehensive set of representative baselines, the global graph embeddings of the filtered background states are subjected to K-means clustering. This partitions the continuous trajectories of the opposing class into distinct latent micro-states, effectively capturing its accessible conformational landscape without redundancy.
3. *Medoid Extraction:* From each discovered cluster, the most representative temporal window (the medoid) is extracted to serve as a discrete physical baseline for the IG calculation. In this work, we consistently selected N=5 medoids per background class, as this number provides an optimal balance between representing the full structural diversity of the opposing state and maintaining computational tractability (**Extended Data Fig. 8**).
4. *Topology Unification:* Because sliding windows exhibit varying dynamic contact topologies, we constructed a unified interaction network. If a specific inter-residue contact exists in the target graph but is absent in a medoid baseline (or vice versa), it is aligned via the injection of a “ghost edge”. Initialized with zero-intensity RBF features, these ghost edges mathematically represent interactions at or beyond the spatial cutoff, allowing the IG algorithm to evaluate transient contacts continuously.
5. *Expected Attribution:* The final structural attribution is computed as the expectation value of the integrated gradients across these multiple physical baselines, weighted by the fractional population of each corresponding K-means cluster.

#### Spatiotemporal Aggregation

To translate edge-level interactions into intuitive, per-residue measures, the attribution score of each edge in a frame is distributed equally between its two connecting nodes. These instantaneous scores are then summed across the temporal dimension of the sliding window, yielding a unified importance score that reflects the dynamic role of each residue.

#### Cross-Validation Consensus and 3D Mapping

To extract a generalized structural consensus and eliminate trajectory-specific noise, attribution scores are aggregated across all Leave-One-Replica-Out cross-validation folds. For each residue, we compute the mean of its window-aggregated importance scores. This mean robustly captures the average contribution of a residue across the conformational ensemble. These scores are finally mapped onto a reference 3D structure to visualize the critical regions defining the conformational ensembles.

### Molecular Dynamics Simulations

#### T4-Lysozyme and PDZ3 Domain Systems

Molecular dynamics (MD) simulations for T4-Lysozyme (wild-type and L99A mutant) and the PDZ3 domain (apo and holo states) were performed using the AMBER software package. Starting conformations were generated by mutating the WT conformation for T4-Lysozyme and removing the peptide from the holo structure for PDZ3 domain. Using a common reference conformation and allowing the system to relax into distinct conformational states helps prevent the learning task from becoming trivial: different starting geometries for the different states would have provided the model with a strong initial geometric bias. For each system, initial coordinates were solvated in a periodic box of explicit water molecules and neutralized by adding counterions (Na+, Cl-) to reach physiological concentration. To ensure proper system stabilization prior to production runs, a rigorous multi-step preparation protocol was employed. Initially, the solvent and ions were minimized while the solute (protein and peptide, where applicable) was held in place using harmonic restraints with a force constant of 100 kcal/mol/Å^2^. This initial phase consisted of 2,000 minimization cycles (1,000 steps of steepest descent followed by conjugate gradient), deliberately omitting the SHAKE algorithm to allow for the resolution of local steric clashes. Subsequently, the system was gradually heated from 100 K to 298 K in the NVT ensemble over 1 ns (using a 1 fs integration time step), regulated by a Langevin thermostat with a collision frequency of 1.0 ps^−1^. During the heating phase, a 10 kcal/mol/Å^2^ restraint was applied to the protein, and the SHAKE algorithm was activated to constrain bonds involving hydrogen atoms. Following heating, the systems were subjected to a comprehensive equilibration phase in the isothermal-isobaric (NPT) ensemble at a reference pressure of 1 atm, controlled by a Monte Carlo barostat. This phase involved the gradual, stepwise release of harmonic restraints on the protein backbone atoms through successive runs of 1 ns, 2 ns, and further cycles totaling approximately 10 ns, progressively reducing the force constant from 10.0 to 0.2 kcal/mol/Å^2^. Midway through the equilibration protocol, a brief secondary minimization phase (1,000 cycles) was introduced to facilitate side-chain repacking. Production simulations were performed in the NPT ensemble at 300 K using a 2 fs time step. Long-range electrostatic interactions were computed using the Particle Mesh Ewald (PME) method, with a non-bonded interaction cutoff set to 10.0 Å. To guarantee exhaustive and statistically robust conformational sampling, multiple independent replicas were generated by assigning randomized initial velocities. Specifically, for T4-Lysozyme, five independent 500 ns replicas were produced for each target state. For the PDZ3 domain, to adequately capture slower allosteric motions, five independent 1 µs replicas were generated for both the apo and holo states. *Adenosine A2A GPCRmd Datasets*

To evaluate the extraction of activation signatures within transmembrane G-protein-coupled receptors, molecular dynamics trajectories of the Adenosine A2A receptor were retrieved from the GPCRmd database^41^ (http://gpcrmd.org). For the classification task, independent trajectories of the receptor bound to four distinct agonist ligands (defining the active state) and four distinct antagonist ligands (defining the inactive state) were selected. The specific complexes utilized are detailed in Table 2.

**Table 2.**
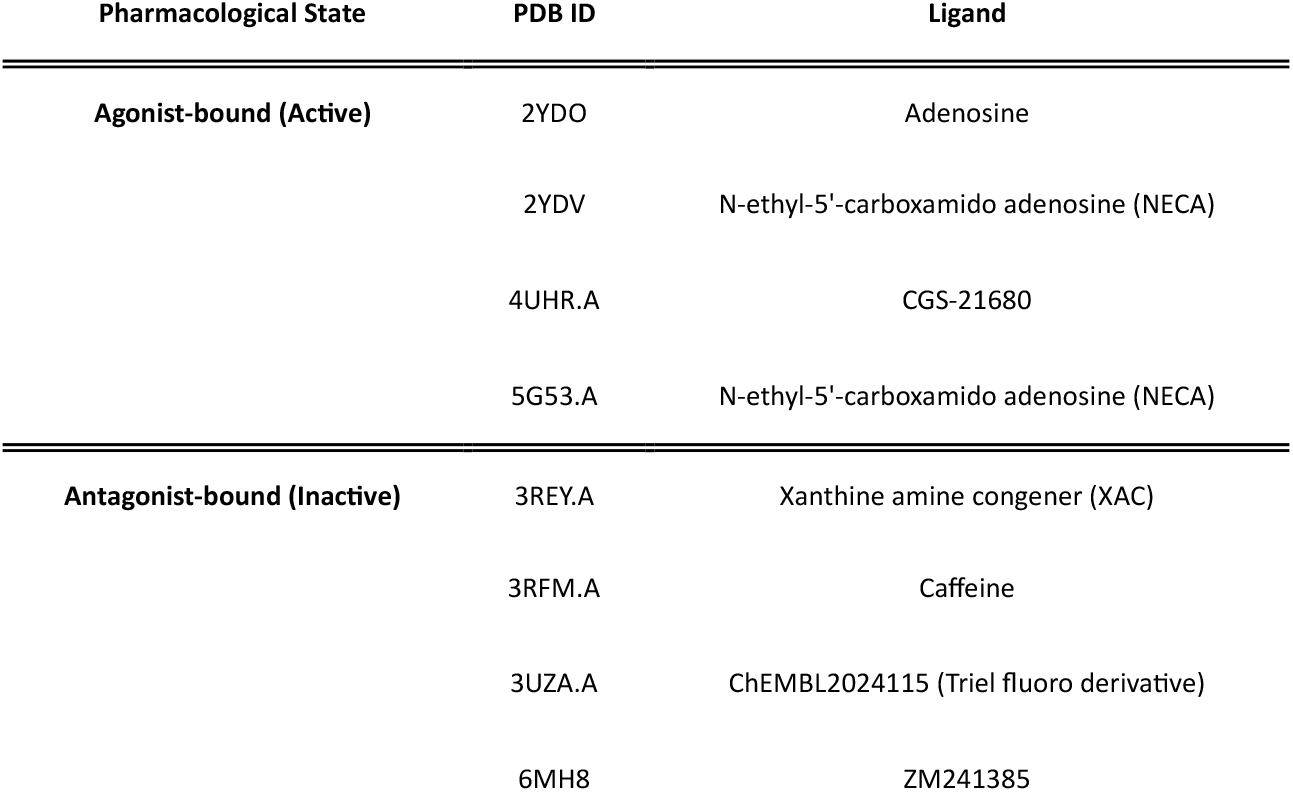
Adenosine A2A receptor simulation datasets. Summary of the GPCRmd trajectories utilized for the extraction of universal activation signatures. The table details the pharmacological state, original PDB identifiers, and specific ligands for the agonist-bound (Active) and antagonist-bound (Inactive) complexes.

| Pharmacological State | PDB ID | Ligand |
| --- | --- | --- |
| Agonist-bound (Active) | 2YDO | Adenosine |
|  | 2YDV | N-ethyl-5'-carboxamido adenosine (NECA) |
|  | 4UHR.A | CGS-21680 |
|  | 5G53.A | N-ethyl-5'-carboxamido adenosine (NECA) |
| Antagonist-bound (Inactive) | 3REY.A | Xanthine amine congener (XAC) |
|  | 3RFM.A | Caffeine |
|  | 3UZA.A | ChEMBL2024115 (Triel fluoro derivative) |
|  | 6MH8 | ZM241385 |

For each ligand-receptor complex, three independent simulation replicas of approximately 500 ns each were analyzed. As described in the main text, to evaluate structural dynamics independent of initial coordinate biases, the agonist-bound simulations were additionally compared against corresponding active-apo trajectories available in the GPCRmd repository. These apo trajectories were simulated starting from the identical active conformational coordinates as the holo simulations, following the *in silico* removal of the ligand prior to the execution of the simulation protocol.

## Acknowledgement

We acknowledge CINECA for the availability of high-performance computing resources as part of the agreement with the University of Milano-Bicocca and the award under the ISCRA initiative (project HP10BBT0CP).

## Code availability

The code to train GISTnet-MD and perform the explainable AI pipeline is freely available on GitHub at https://github.com/MottaStefano/GISTnet-MD.

## Author Contributions

S.Mo. conceived the idea, developed the methodology, wrote the software, led the formal analysis, and curated the data. S.Mo., A.P., and M.M. conceptualized the broader study and discussed the results. A.P. and M.M. jointly supervised the work. M.M. contributed to the validation of the methodology, G.S. contributed to performing standard formal analysis. S.Mo. wrote the original draft. F.H.N., A.P., M.M., and S.Ma. revised the manuscript. All authors approved the submitted manuscript and agreed to be personally accountable for their own contributions to the work.

