## Extended Data Figures for "Spatiotemporal graph neural networks reveal conformational binding signature in protein dynamics"

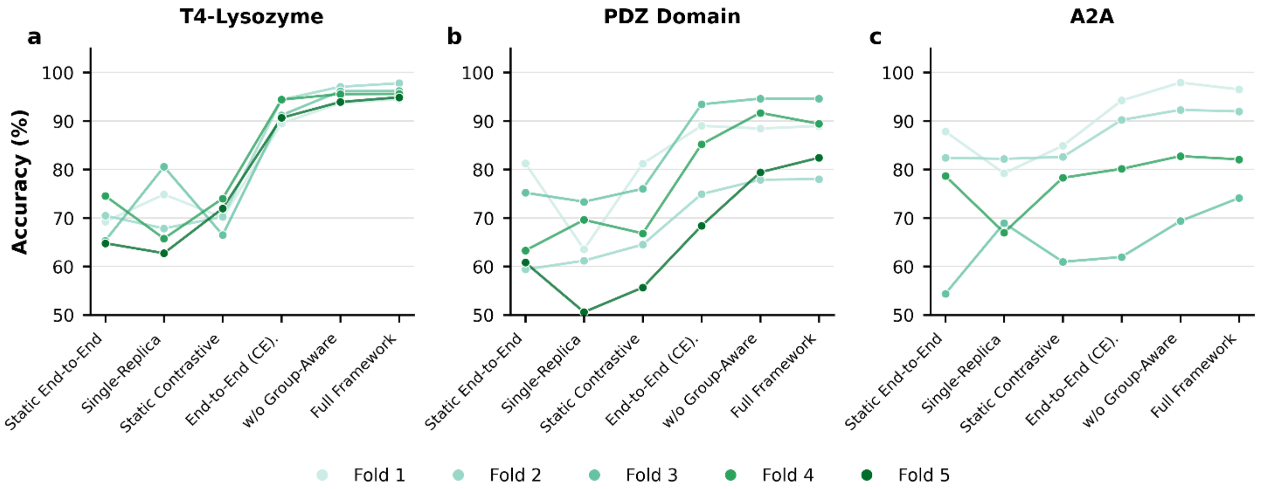


**Extended Data Fig. 1 | Trajectory analysis of individual cross-validation folds.** Slopegraphs tracking the validation accuracy of independent data splits (folds) across six model configurations for **a**, T4-Lysozyme; **b**, PDZ Domain; and **c**, A2A Receptor. Each coloured line represents one held-out validation group, corresponding to an independent MD replica for T4 lysozyme and PDZ3, or a ligand-defined group for A2A. Data points denote the mean accuracy averaged over three independent random seeds. The plots show how performance changes across architectural variants and reveal fold-specific variability that may be obscured in aggregate bar plots.

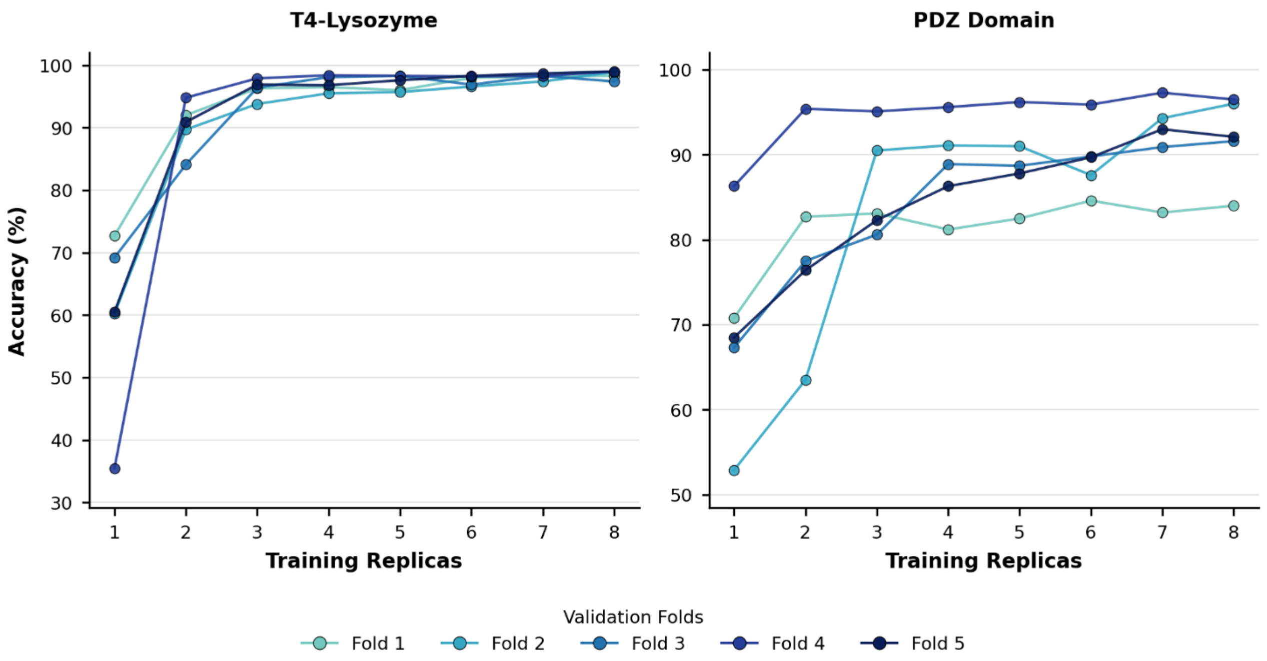


**Extended Data Fig. 2 | Convergence trajectories across individual cross-validation folds.** Line charts tracking the validation accuracy of independent data splits (folds) as a function of training set size for **a**, T4-Lysozyme and **b**, PDZ Domain. Each colored line traces the trajectory of a single cross-validation fold, illustrating the performance gain as the number of training MD replicas increases from 1 to 8. Data points represent the accuracy achieved on a specific validation fold (including two replicas for each state).The overall upward trends show that adding independent training replicas improves generalisation to unseen trajectories, while the fold-to-fold variability reflects system-specific differences in sampling difficulty. T4 lysozyme reaches stable high accuracy with fewer training replicas, whereas PDZ3 requires additional replicas to achieve comparable stability.

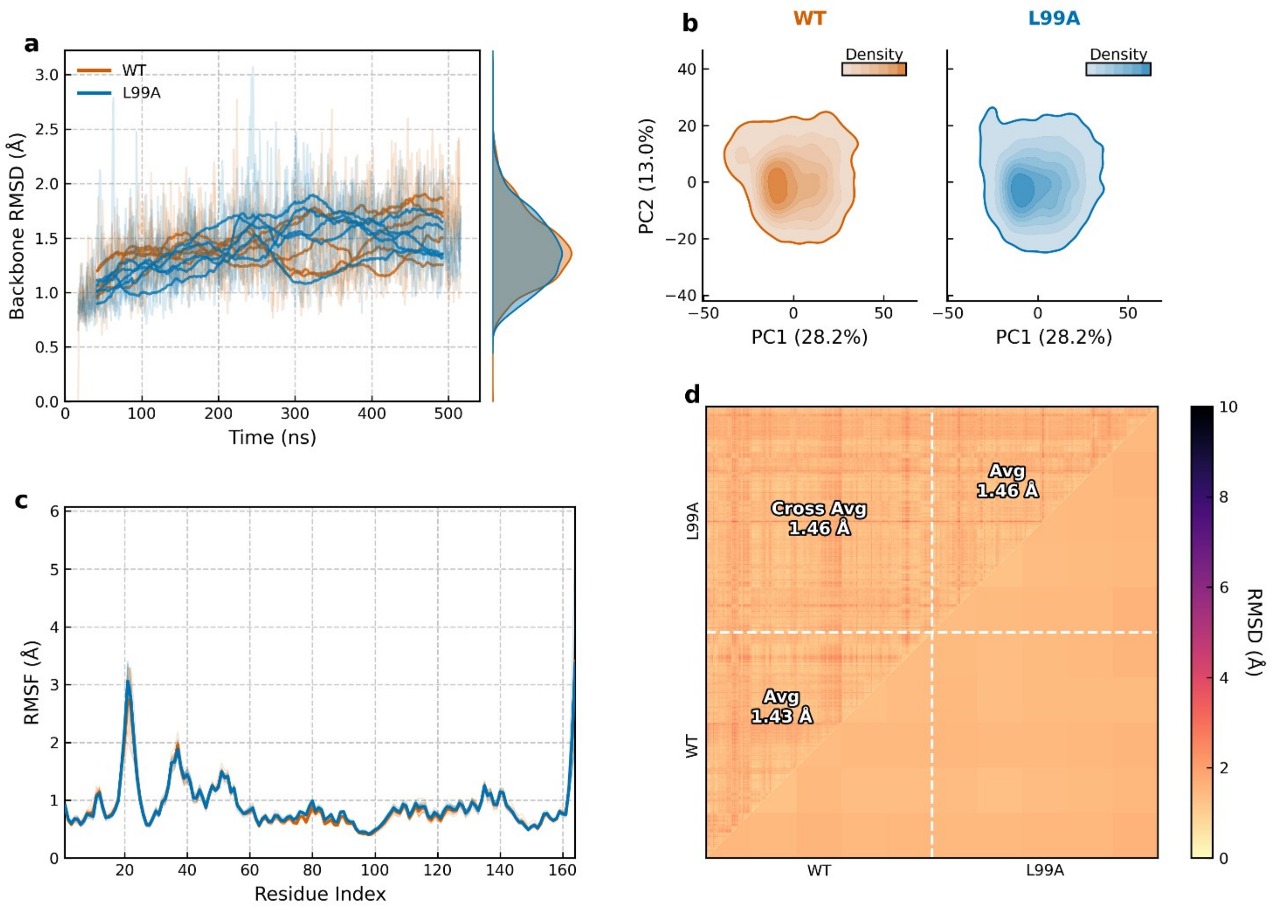


**Extended Data Fig. 3 | Standard molecular dynamics characterization of T4-Lysozyme wild-type and L99A conformational ensembles. a,** Time evolution of the backbone Root Mean Square Deviation (RMSD) calculated over five independent 500 ns simulation replicas for both the wild-type (WT, orange) and the cavity-forming L99A mutant (blue). Faded lines depict the raw data, while solid lines represent a 50-frame moving average. The right-hand panel displays the corresponding Kernel Density Estimate (KDE) distributions, highlighting the extensive overlap in the global structural deviation of the two macroscopic states. **b,** Principal Component Analysis (PCA) projection of the backbone coordinates into a shared essential subspace. The 2D density contour plots demonstrate that WT and L99A populate highly overlapping regions of the projected conformational landscape along the first two principal components (PC1 and PC2, capturing 28.2% and 13.0% of the total variance, respectively). **c,** Per-residue Root Mean Square Fluctuation (RMSF) profiles. Solid lines indicate the mean RMSF across the five independent replicas, with shaded regions representing the standard deviation. Both ensembles exhibit nearly identical local flexibility profiles across the entire sequence, with slight variability in the region comprised between residues 70-90. **d,** All-vs-all 2D RMSD matrix across the concatenated trajectories (subsampled for visualization). The lower triangle displays the frame-by-frame pairwise RMSD, while the upper triangle reports the block-averaged RMSD for each specific replica-to-replica comparison. The global intra-ensemble averages (1.43 Å for WT vs. WT; 1.46 Å for L99A vs. L99A) are virtually indistinguishable from the cross-ensemble average (1.46 Å).

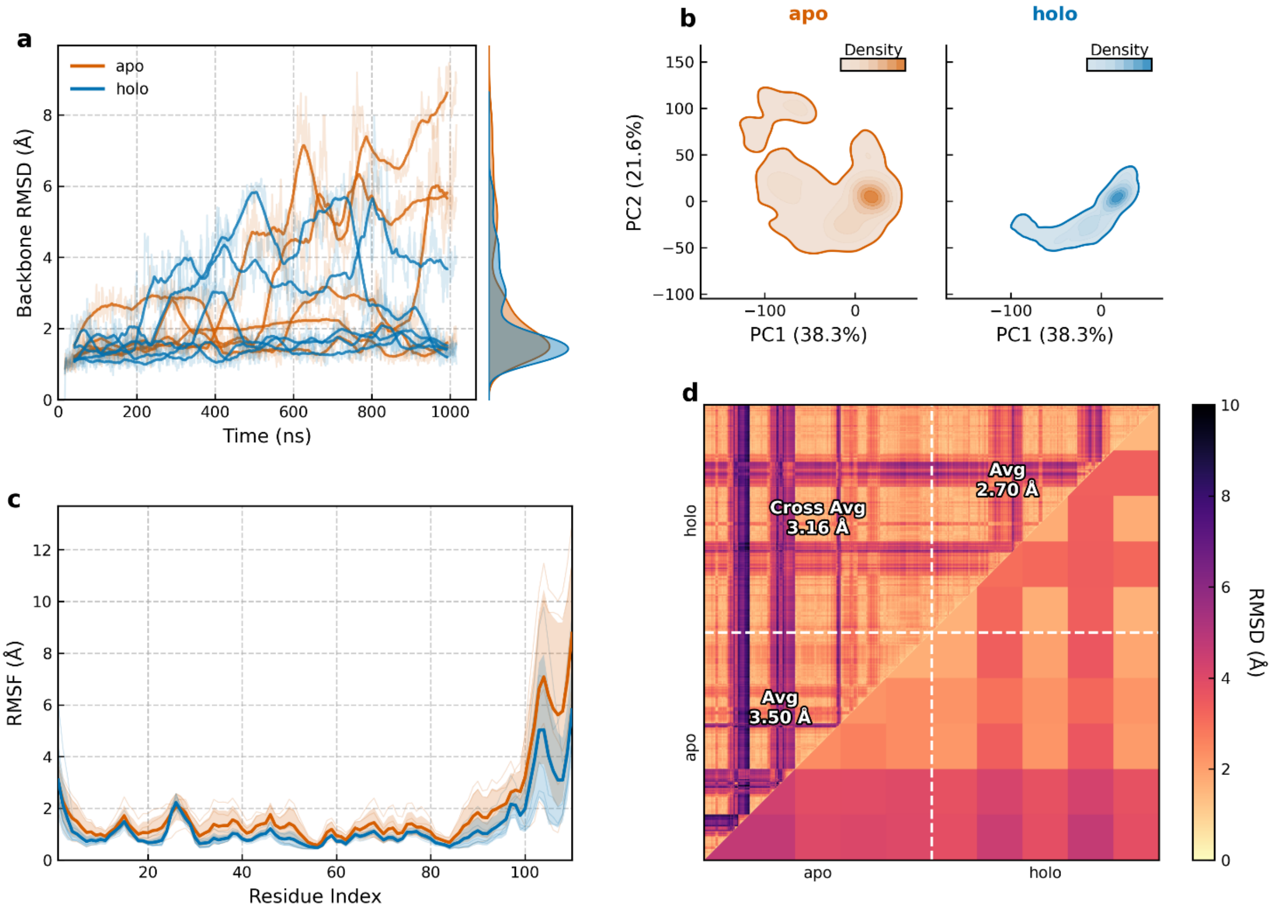


**Extended Data Fig. 4 | Standard molecular dynamics characterization of the PDZ3 domain apo and holo (peptide-bound) conformational ensembles. a,** Time evolution of the backbone Root Mean Square Deviation (RMSD) calculated over five independent 1 µs simulation replicas for the apo state (orange) and the holo state (blue). Faded lines depict the raw data, while solid lines represent a 50-frame moving average. The right-hand panel displays the corresponding Kernel Density Estimate (KDE) distributions. Notably, both states exhibit pronounced inter-replica divergence and massive structural fluctuations (reaching >8 Å), indicative of a highly dynamic and flexible ensemble. **b,** Principal Component Analysis (PCA) projection of the backbone coordinates into a shared essential subspace. The 2D density contour plots strikingly illustrate the thermodynamic discrepancy between the two states. The unliganded apo ensemble explores a vast, highly dispersed conformational landscape comprising multiple metastable basins. Conversely, peptide recognition funnels this intrinsic flexibility, restricting the holo domain to a narrow, conformational state along the first two principal components (PC1: 38.3%, PC2: 21.6%). **c,** Per-residue Root Mean Square Fluctuation (RMSF) profiles. Solid lines indicate the mean RMSF across the five independent replicas, with shaded regions representing the standard deviation. The apo state exhibits substantially higher local flexibility, most prominently at the C-terminal region (residues 100-110) and specific loop segments. Peptide binding dampens these large-scale fluctuations globally, providing a classic signature associated with of distributed dynamic allostery. **d,** All-vs-all 2D RMSD matrix across the concatenated trajectories (subsampled for visualization). The strong block-diagonal patterns and the high global intra-ensemble average for the apo state (3.50 Å) quantitatively confirm the presence of highly divergent, replica-specific conformational exploration. The holo state exhibits higher internal consistency (Avg 2.70 Å). Collectively, the extreme variance, the stochastic inter-replica divergence, and the highly distributed nature of these dynamic changes emphasize that standard global metrics do not directly resolve subtle, residue-level dynamic signatures, reinforcing the necessity for the robust, noise-filtering spatiotemporal feature extraction provided by GISTnet-MD.

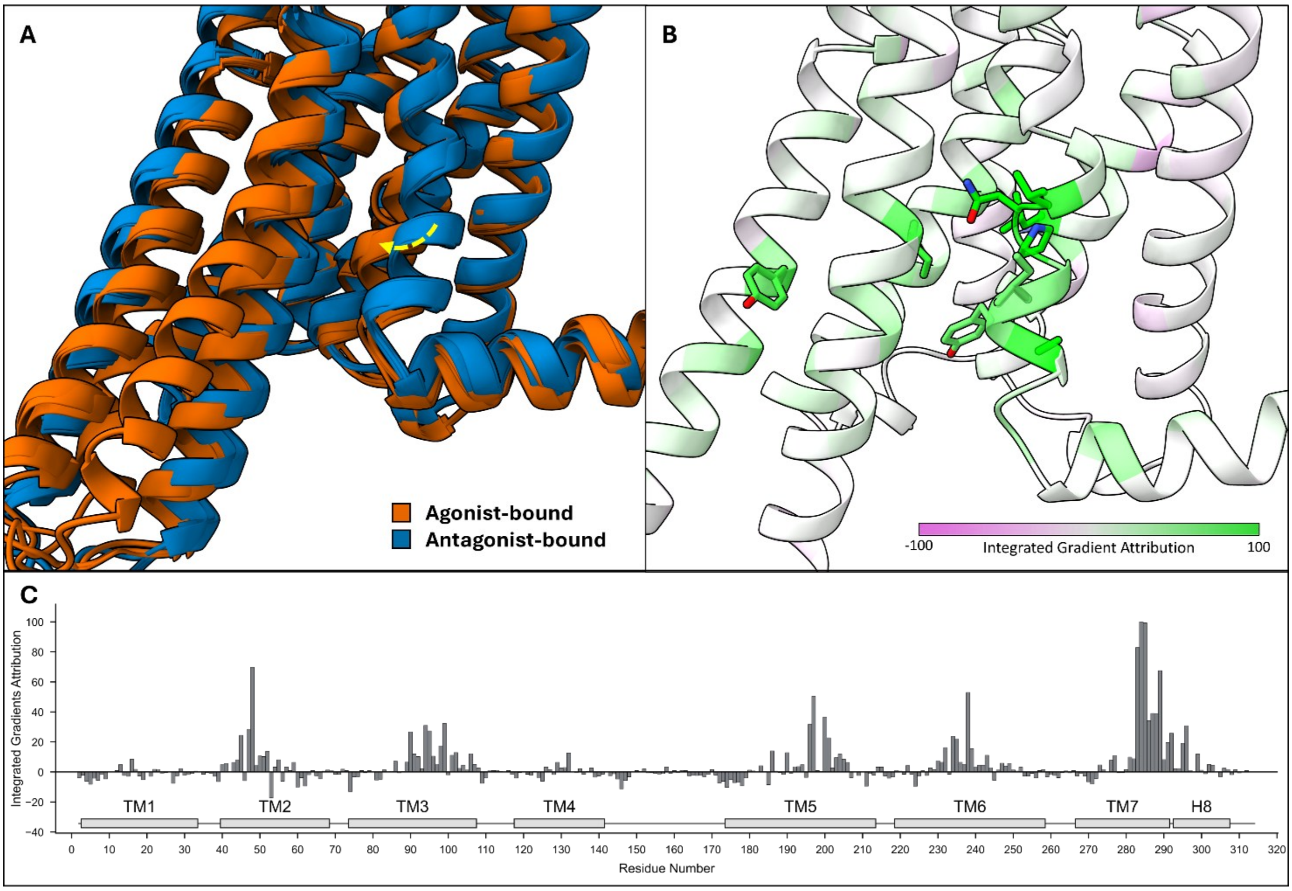


**Extended Data Fig. 5 | Network identification of structural biases in the Agonist vs Antagonist classification.** **a**, Structural superposition of the initial coordinates utilized for the molecular dynamics simulations of the agonist-bound (active state, orange) and antagonist-bound (inactive state, blue) A2A receptor ensembles. The yellow dashed arrow highlights the macroscopic, pre-existing inward shift of the intracellular half of Transmembrane Helix 7 (TM7), encompassing the canonical NPxxY microswitch. **b**, 3D structural mapping of the IG scores on the A2A structure, with the primary classification-driving residues colored in green. **c**, 1D consensus IG profile for the full, unmasked GISTnet-MD model trained to differentiate these two states. Accurately reflecting the geometric disparities of the starting structures, the network's classification demonstrates a strong reliance on this pre-existing structural difference, assigning dominant attribution scores to the divergent TM7 region (residues 280-290).

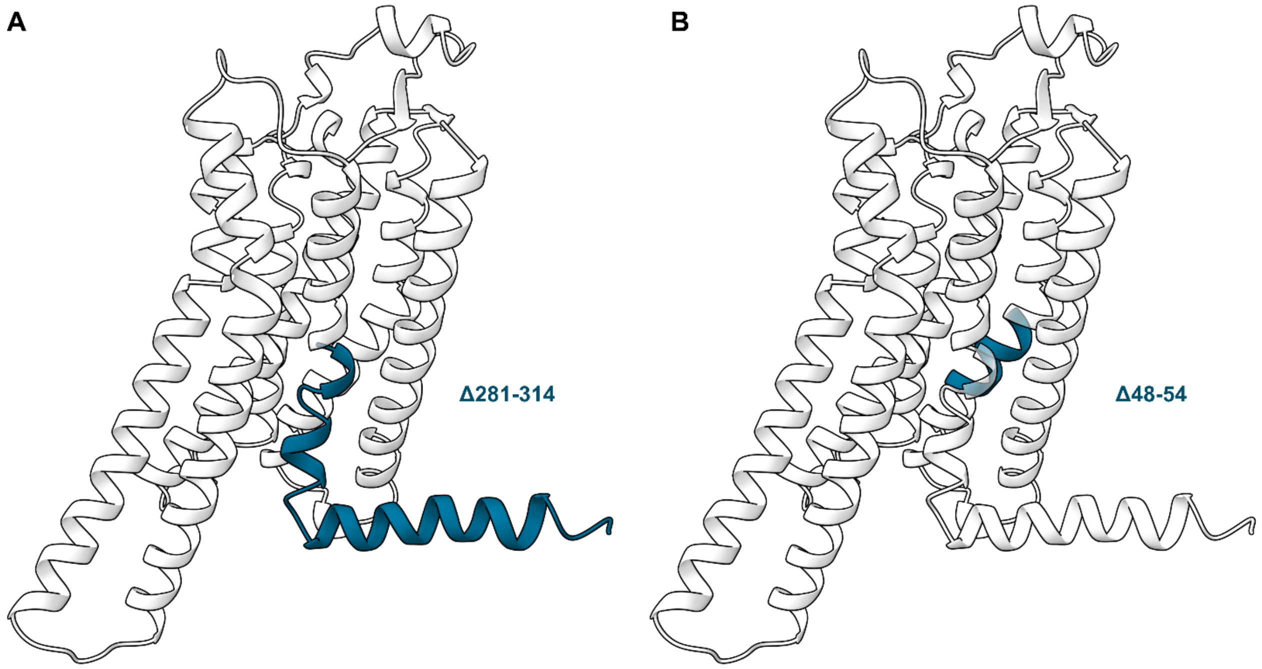

**Extended Data Fig. 6 | Structural mapping of the regions masked during the in silico ablation experiments on the Adenosine A2A receptor. A**, Three-dimensional representation of the A2A receptor highlighting the intracellular segment encompassing the lower portion of Transmembrane Helix 7 (TM7) and the adjacent C-terminal region (Δ281–314, highlighted in blue). This region was entirely removed from the input graph to train the truncated model for the agonist-bound versus antagonist-bound classification task. **B**, Structure of the A2A receptor illustrating the localized sodium-binding pocket (Δ48–54, highlighted in blue). This specific segment was masked during the active-holo versus active-apo classification task to mathematically decouple the dynamic allosteric analysis from the dominant electrostatic signal of the D52 protonation state.

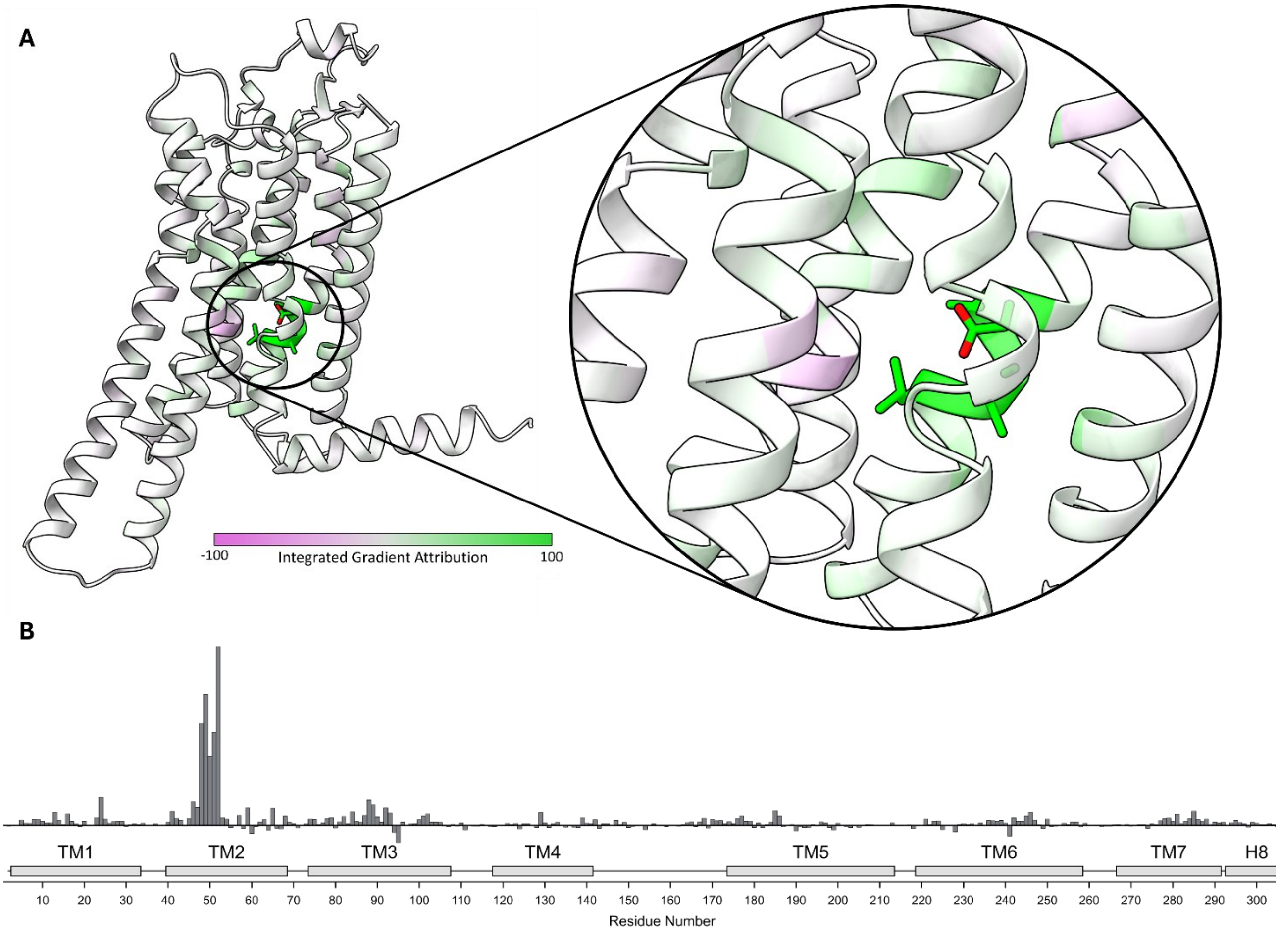


**Extended Data Fig. 7 | Network identification of localized electrostatic bias in the unmasked Agonist vs Apo classification. A**, 3D structural mapping of the consensus Integrated Gradients (IG) scores on the A2A receptor, with a magnified inset detailing the deeply buried sodium pocket. The primary classification-driving residue, Asp52, is shown in stick representation and prominently colored in green. **B**, 1D consensus IG profile for the full, unmasked GISTnet-MD model trained to differentiate the agonist-bound (active-holo) and active-apo ensembles originating from identical coordinates. Lacking macroscopic structural differences, the network entirely bypasses the subtle transmembrane allostery, latching instead onto the massive discrepancy generated by the differential protonation state of Asp52 (protonated in holo, deprotonated in apo).

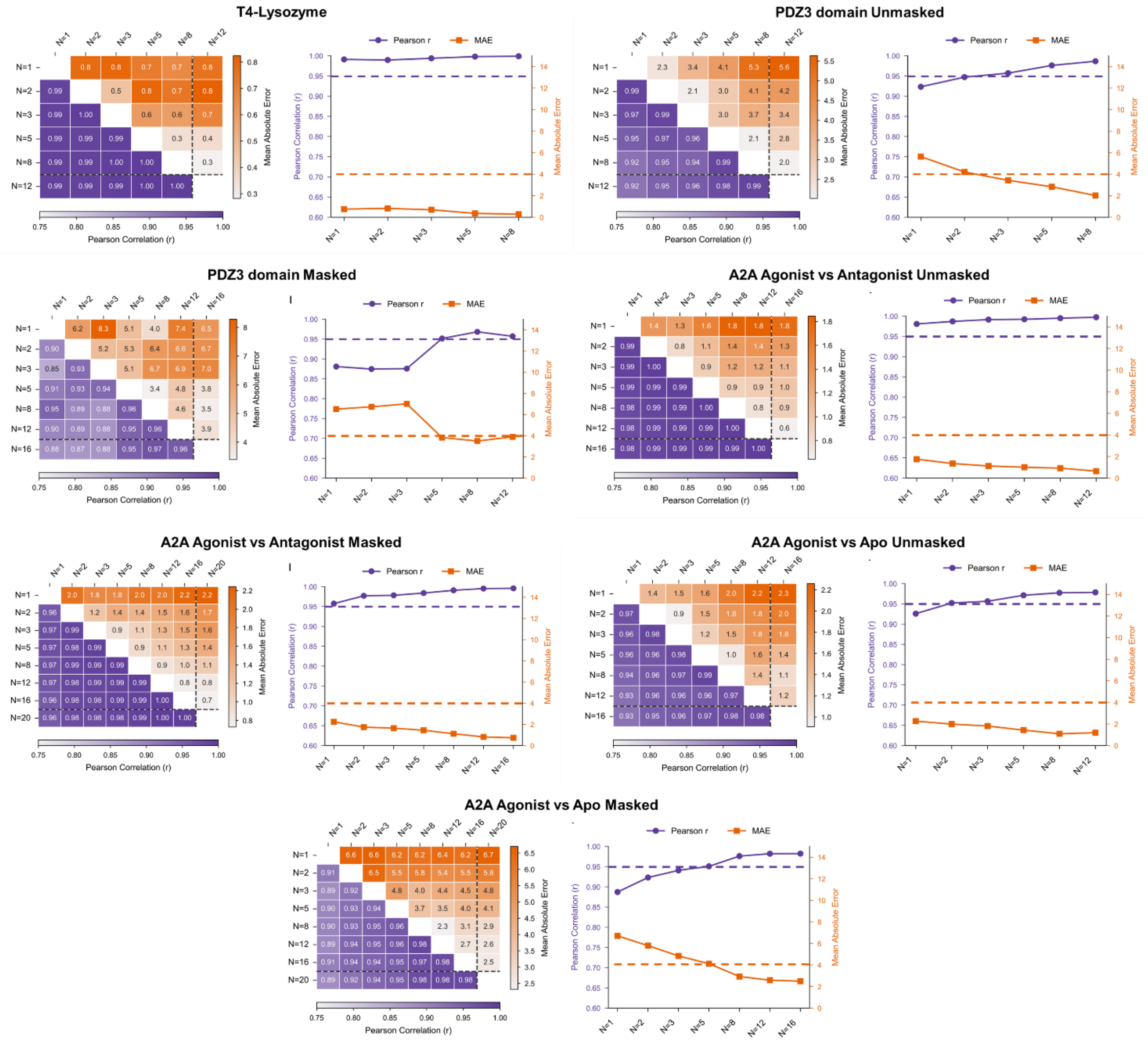


**Extended Data Fig. 8 | Convergence analysis of the structural attribution baseline across the evaluated molecular systems. a–g**, Stability of the Integrated Gradients (IG) attribution profiles as a function of the number of representative background windows (*N*) used to construct the baseline. Background windows were sampled from the contrasting thermodynamic ensemble using K-means clustering of the latent embeddings, and medoid windows from an increasing number of clusters (*N*) were evaluated against the highest-*N* reference baseline. The panels detail the analysis for the specific comparative tasks: T4-Lysozyme (**a**), unmasked PDZ3 domain (**b**), masked PDZ3 domain (**c**), unmasked A2A Agonist vs. Antagonist (**d**), masked A2A Agonist vs. Antagonist (**e**), unmasked A2A Agonist vs. Apo (**f**), and masked A2A Agonist vs. Apo (**g**). Within each panel, the left sub-plot presents a hybrid matrix quantifying the pairwise agreement between baseline parameterizations. The lower-left triangle (purple gradient) reports the Pearson correlation coefficient (*r*), while the upper-right triangle (orange gradient) reports the Mean Absolute Error (MAE). The reference baseline (higher number of *N* investigated) is spatially isolated by a black dashed line. The right sub-plot tracks the progression of these distance metrics between each configuration and the reference baseline as a function of the sample size. Purple circles (left y-axis) indicate the Pearson correlation (*r*), and orange squares (right y-axis) denote the MAE. Overall, the convergence of both metrics supports the selected baseline size (*N=5*) as a practical balance between attribution stability and computational cost, while indicating that more complex masked tasks may benefit from denser baseline sampling.
