## Supplementary Notes for "Spatiotemporal graph neural networks reveal conformational binding signature in protein dynamics"

### Supplementary Note 1: Dissecting the 1D-CNN via Temporal Data Shuffling

To investigate the specific features extracted by the 1D-CNN from the sliding temporal windows, a series of temporal shuffling experiments was performed (**Supplementary Fig. 1**). Two distinct perturbation strategies were applied: *window shuffling* (randomizing the chronological order of frames strictly within a given temporal window) and *global shuffling* (constructing windows by randomly sampling frames from the entire global trajectory). These perturbations were applied asymmetrically: exclusively during training (train), exclusively during validation (val), or during both phases (all).

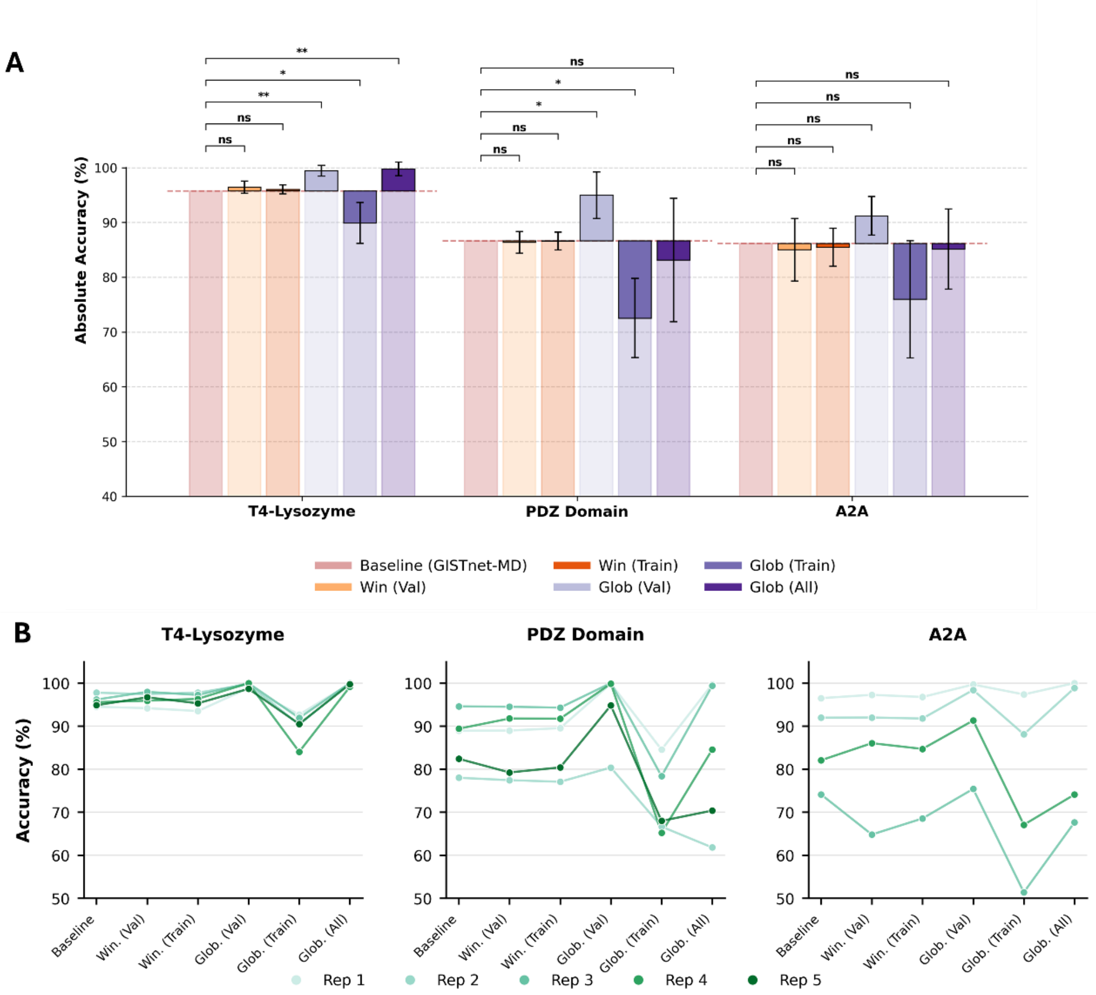


**Supplementary Fig. 1 | Impact of temporal data shuffling on model performance.** Evaluation of the GISTnet-MD architecture under different temporal perturbation strategies across the three evaluated protein systems. **A**, Bar plots displaying the validation accuracy of the unperturbed baseline compared to models where temporal correlations were disrupted via window-based or global shuffling, applied exclusively during validation, exclusively during training, or during both phases (All). Semi-transparent bars indicate the absolute mean classification accuracy across independent cross-validation folds (*n=5*). Solid colored bars highlight the performance gap, extending vertically from the baseline reference (horizontal dashed line) to the shuffled variant's accuracy, accommodating both performance degradation and regularization effects. Error bars denote the standard deviation of the paired, per-replica performance differences, isolating the impact of the temporal perturbation from trajectory-specific noise. Statistical significance was determined using two-sided paired t-tests against the baseline configuration (* $P < 0.05$, ** $P < 0.01$, *** $P < 0.001$, ns = not significant). **B**, Slopegraphs tracking the validation accuracy of independent data splits (folds) under the progressive temporal perturbation scenarios. Each colored line traces the trajectory of a single cross-validation replica to assess fold-specific sensitivity.

Window shuffling exhibited minimal impact on classification accuracy across all evaluated systems. This observation indicates that the 1D-CNN does not memorize rigid chronological sequences. Instead, the module functions as a dynamic temporal filter, weighting the relative contributions of the conformations sampled within the localized timeframe and evaluating the window as a localized equilibrium ensemble, independent of presentation order. Conversely, global shuffling yielded substantial performance shifts. When applied exclusively to the validation replica, accuracy increased. When applied exclusively during training, accuracy declined significantly. When applied to both phases, accuracy improved for the T4-Lysozyme system but deteriorated or remained unchanged with increased variance for the structurally more complex systems (PDZ domain and A2A receptor).

This behavior can be explained by considering the conformational variance within the temporal windows relative to the protein's energy landscape. A chronologically ordered window represents a continuous trajectory within a specific local energy basin, characterized by low structural variance. The model is therefore required to learn local dynamic features, such as specific structural fluctuation patterns. Validating such a model on globally shuffled data artificially constructs windows containing highly diverse conformations sampled from the entire global landscape. This exposes the network simultaneously to representative macroscopic states, trivially simplifying the classification task and generating the observed accuracy increase in the validation shuffling scenario as a diagnostic artifact of artificial data mixing. Correspondingly, training on globally shuffled windows forces the network to rely on these macroscopic structural outliers. The model learns to identify large-scale geometric differences but fails to develop the sensitivity required to analyze subtle local dynamic variations. Consequently, when validated on standard, chronologically ordered windows (trapped in a single local basin), the model struggles to assign the state confidently due to a severe distribution mismatch between the training and validation data. Furthermore, this dynamic explains the discrepancy between the simple T4-Lysozyme system and the complex biological systems (PDZ, A2A) in the "all" scenario. In T4-Lysozyme, where the trajectory cleanly populates a single macrostate, global shuffling facilitates classification. In realistic biological simulations, which often sample transient confounding macrostates, global shuffling distributes frames from these detached states across the entire validation set. This systematic dilution of the temporal windows across the trajectory explains the performance degradation and increased cross-replica variance observed in the complex systems. A detailed benchmarking of the computational cost per cross-validation fold under different temporal configurations is provided in **Supplementary Table 1**.

**Supplementary Table 1:** Per CV-fold computational cost of calculations with different temporal hyperparameters. Results are just an indication because they may vary depending on the number of training epochs performed (patient may stop calculations earlier) and speed may be optimized acting on the microbatch size. The frame number is the total number of frames for the training set.

| System | Window size | offset | Stride (ps) | Residue number | Frame Number | Computational cost per CV-fold (min) |
|---|---|---|---|---|---|---|
| T4-Lysozyme | 5 | 1 | 100 | 164 | 50,000 | 32 |
| T4-Lysozyme | 25 | 10 | 100 | 164 | 50,000 | 34 |
| T4-Lysozyme | 25 | 1 | 100 | 164 | 50,000 | 135 |
| T4-Lysozyme | 100 | 1 | 100 | 164 | 50,000 | 1048 |
| PDZ | 5 | 1 | 100 | 110 | 100,000 | 52 |
| PDZ | 25 | 10 | 100 | 110 | 100,000 | 65 |
| PDZ | 25 | 1 | 100 | 110 | 100,000 | 294 |
| PDZ | 100 | 1 | 100 | 110 | 100,000 | 1101 |
| A2A | 5 | 1 | 200 | 313 | 60,000 | 97 |
| A2A | 25 | 10 | 200 | 313 | 60,000 | 138 |
| A2A | 25 | 1 | 200 | 313 | 60,000 | 519 |
| A2A | 100 | 1 | 200 | 313 | 60,000 | 2750 |

### Supplementary Note 2: Geometric Analysis of the T4-Lysozyme L99A Cavity

The L99A substitution in T4-Lysozyme is well-characterized for generating a buried, hydrophobic cavity that binds various small non-polar ligands, shifting the conformational ensemble toward a binding-ready state. To quantify the internal volume, we implemented a custom Monte Carlo integration algorithm applied to each analyzed frame of the MD trajectories. For each analyzed frame, a bounding sphere with a radius of 8.0 Å was defined. This sphere was centered on the center of geometry of a specific set of pocket-lining and adjacent residues (residues 95, 98, 99, 111, 117, 118, 121, 126, 129, 133, 149, 150, and 153). An ensemble of 200,000 points was uniformly generated within this bounding sphere. To evaluate the accessible pocket volume, points were designated as occupied, and thus excluded from the free volume, if located within the Van der Waals radius of any adjacent protein atom plus a probe radius of 1.0 Å. To restrict the volume calculation to the primary contiguous binding pocket and exclude isolated interstitial voids, a graph-based spatial clustering step was applied. A spatial KD-Tree was constructed for all free points, and pairs within a 0.5 Å distance threshold were connected to generate a sparse adjacency matrix. Following the extraction of connected components, exclusively the points belonging to the largest contiguous spatial cluster were retained. The accessible cavity volume was computed as the fraction of retained free points relative to the total generated points, multiplied by the analytical volume of the bounding sphere.

As shown in **Supplementary Fig. 2a**, the time evolution of the cavity volume demonstrates a sustained expansion in the L99A mutant compared to the WT. While both ensembles exhibit rapid thermal fluctuations, the mutant maintains a consistently larger internal void throughout the 500 ns simulated timescale. This is further highlighted by the kernel density estimate distributions (**Supplementary Fig. 2b**), which reveal a clear rightward shift in the sampled volumes for the L99A trajectories. Statistical analysis of the replica-averaged volumes confirms a highly significant difference between the two states (**Supplementary Fig. 2c**). Despite this clear geometric distinction at the localized pocket level, standard global geometric metrics such as RMSD, RMSF, and principal component analysis often struggle to robustly differentiate these ensembles due to the subtle and dispersed nature of the structural rearrangements outside the immediate mutation site. This motivates the use of advanced representation learning methods, such as GISTnet-MD, capable of identifying distributed dynamic signatures.

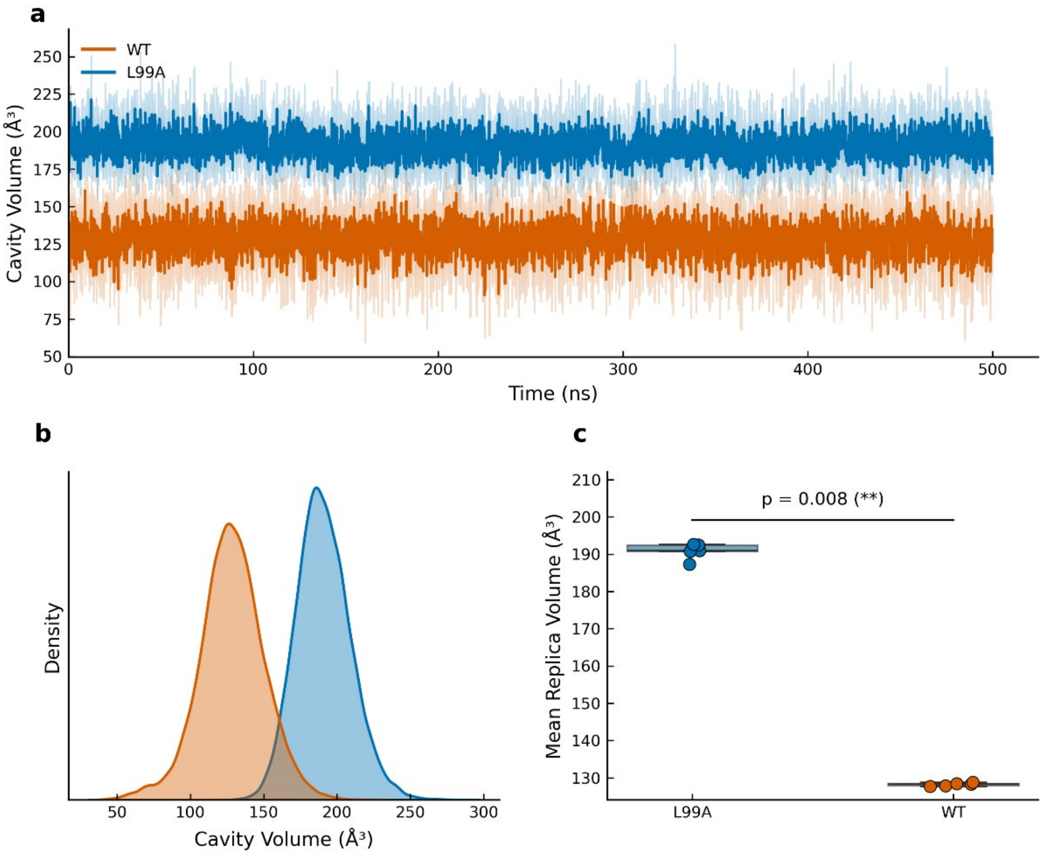

**Supplementary Fig. 2 | Impact of the L99A mutation on the internal cavity volume. a,** Time evolution of the cavity volume over the simulated timescale for the wild-type (WT, orange) and the L99A mutant (blue) ensembles. Solid lines represent the mean volume computed across independent replicas, with shaded areas indicating the 95% confidence intervals. **b,** Kernel density estimate (KDE) distributions of the sampled cavity volumes throughout the trajectories, highlighting a distinct volume expansion induced by the L99A substitution. c, Statistical comparison of the cavity sizes based on replica-averaged volumes. Individual points represent the mean volume of each independent replica, overlaid on box plots. The horizontal bar indicates statistical significance evaluated via a two-sided Mann-Whitney U test (*p = 0.008*, **).

### Supplementary Note 3: Real-time Attribution Tracking of a Transient Unbinding Event

During the analysis of the PDZ3 holo ensemble, we observed a spontaneous partial dissociation of the C-terminal peptide within a specific simulation replica (Replica 2). In this microsecond trajectory, the peptide shifts away from the canonical binding pocket, losing its primary inter-molecular contacts. This rare sampling event provided an opportunity to evaluate the sensitivity of GISTnet-MD to instantaneous structural dynamics and to assess whether the network assigns labels based on the trajectory's provenance or by actively evaluating local biophysical features. Analysis of the model's prediction confidence over time confirms that the spatiotemporal sliding window continuously tracks the physical state of the complex. As shown in **Supplementary Fig. 3a**, prior to the dissociation event, the model robustly classifies the frames as belonging to the holo state. Upon the spatial displacement of the peptide, the classification probability exhibits a sharp transition, reassigning the subsequent temporal windows of this nominally holo trajectory to the apo state.

To extract the mechanistic basis of this reassignment, we evaluated the temporal evolution of the IG attribution scores, projecting them onto the 3D trajectory (**Supplementary Movie 1**). The attribution scores are color-coded to indicate their directional contribution to the classification: positive values (red) indicate spatial regions driving the holo prediction, while negative values (blue) highlight regions actively penalizing the holo classification, pushing the model toward the apo assignment. **Supplementary Fig. 3b** displays the structural attribution of two representative frames extracted from the stable and transient regimes. During the initial phase, characterized by stable peptide binding, the network assigns dense, positive IG attributions (red) to the residues comprising the primary binding cleft (specifically the αB helix and the β2 strand), correctly isolating the dynamic signature of the interaction. Following the partial unbinding event, the physical interaction network is disrupted. Correspondingly, the spatial attribution dynamically updates: the IG scores of the binding site residues invert to negative values (blue), while the surrounding regions exhibit increased signal noise.

This localized inversion of the attribution signal demonstrates that the network's prediction is explicitly coupled to the presence or absence of specific local non-covalent interactions. Rather than failing or generating unstructured noise upon facing a perturbation outside its training distribution, the Explainable AI (xAI) framework actively localizes the precise structural regions where the functional binding signature has been lost. These data confirm that GISTnet-MD operates as an analyzer of instantaneous physical dynamics, capable of resolving continuous thermodynamic state transitions at residue-level resolution.

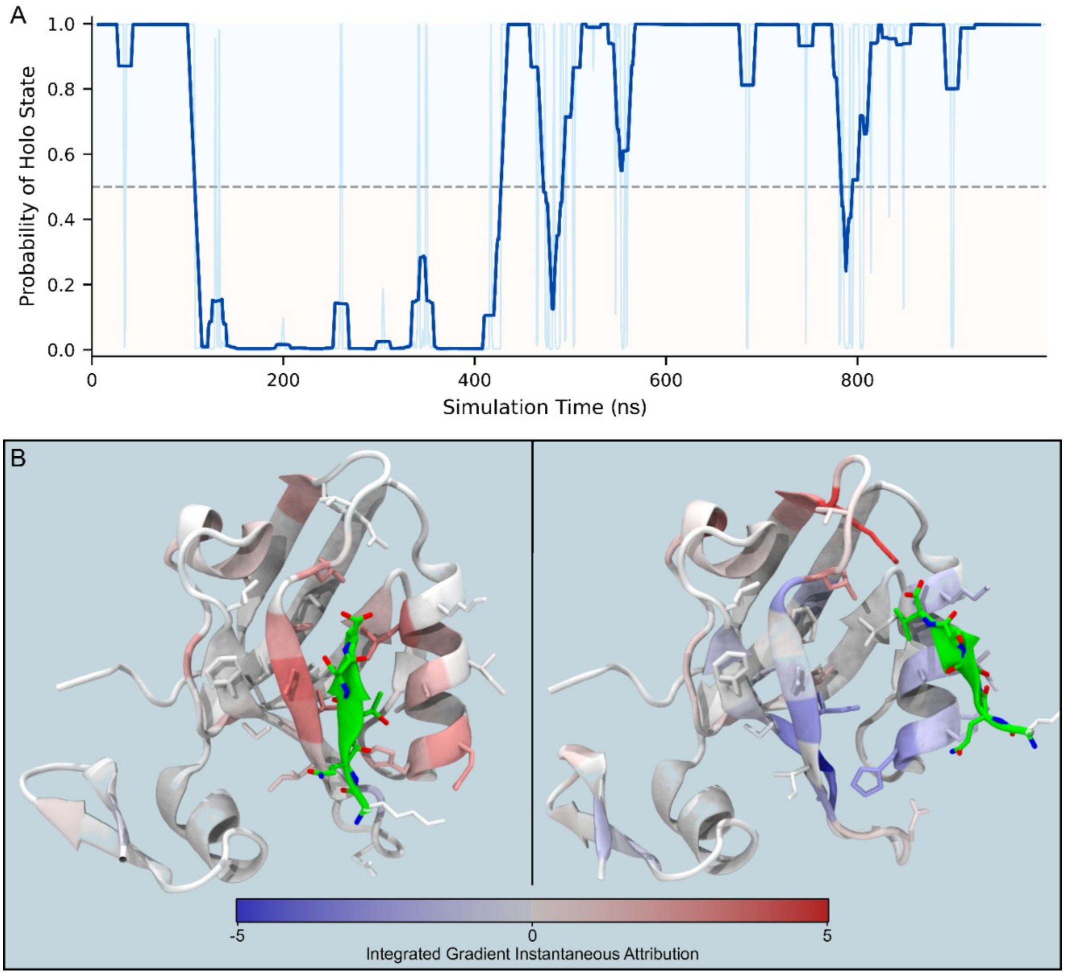


**Supplementary Fig. 3 | Dynamic tracking and structural attribution of a spontaneous partial unbinding event in the PDZ3 domain.** **A**, Time series of the model's prediction confidence (softmax probability) during the PDZ3 holo Replica 2 trajectory. The transition of the prediction from the holo state (peptide-bound) to the apo state coincides with the physical displacement of the peptide. **B**, 3D mapping of the instantaneous IG scores onto representative frames from the simulation. In the stably bound configuration (left panel), the binding site exhibits strong positive attribution. In the partially unbound configuration (right panel), the loss of local interactions drives the attribution scores to negative values at the binding interface.

### Supplementary Note 4: Sensitivity Analysis of Architectural and Temporal Hyperparameters

**Architectural Variations and Feature Configurations:** To systematically evaluate the robustness of GISTnet-MD and determine a highly effective and practical configuration for generalized thermodynamic state identification, a comprehensive sensitivity analysis was performed on key architectural components (**Supplementary Fig. 2**). The analysis was conducted on both the T4-Lysozyme and the PDZ3 Domain (non-masked) systems.

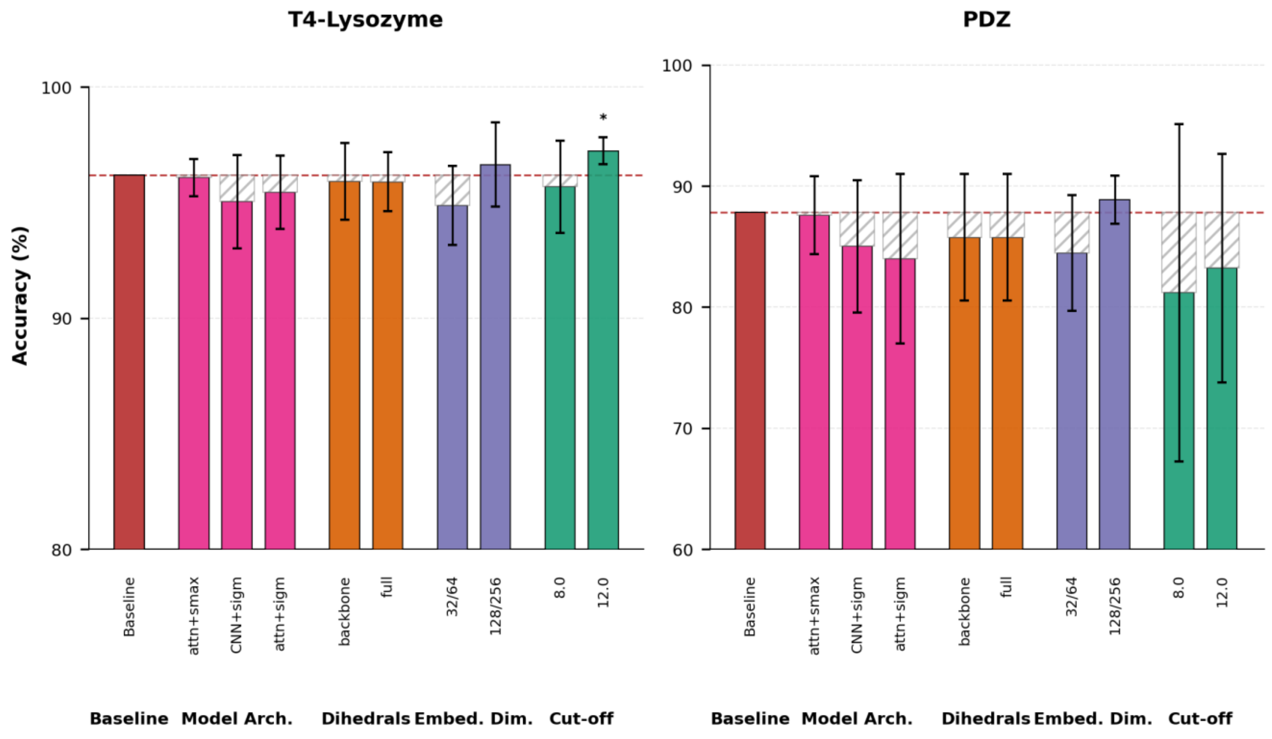


**Supplementary Fig. 4 | Impact of architectural hyperparameters and feature configurations on model accuracy.** Performance is evaluated on two systems: T4-Lysozyme and PDZ Domain. The Baseline configuration (red bar) serves as the baseline. Subsequent bars analyze variations in four categories: model architecture (temporal integration and pooling layers), inclusion of dihedral angle features, node embedding dimension, and interaction cut-off radius. Solid colored bars represent the mean classification accuracy across independent replicas ($n=5$). Hatch-patterned extensions visualize the performance gap when a variant underperforms relative to the baseline. Error bars denote the standard deviation of the paired differences, highlighting the stability of the performance shift. Statistical significance was determined via two-sided paired t-tests against the baseline configuration (* $P < 0.05$, ** $P < 0.01$, *** $P < 0.001$).

The baseline GISTnet-MD architecture utilizes a 1D-Convolutional Neural Network (CNN) for temporal integration and a Softmax activation function within the Global Attention Pooling module. Alternative configurations for these two specific modules were evaluated:

1. *Temporal Integration (CNN vs. Attention): The* baseline CNN extracts local dynamic by applying sliding one-dimensional convolutions along the temporal axis. Alternatively, a Temporal Attention mechanism was implemented, which computes a discrete importance score for each frame within the temporal window, outputting a weighted sum of the residue's embedding history.

2. *Pooling Activation (Softmax vs. Sigmoid):* The baseline Softmax activation enforces a strict sum-to-one constraint across all nodes in the spatial graph, creating a competitive feature selection environment that isolates a sparse set of highly determinant residues. Conversely, the Sigmoid activation evaluates the importance of each node independently, scaling its weight between 0 and 1 without global normalization. This unconstrained approach theoretically accommodates distributed chemical phenomena where multiple nodes may possess high simultaneous relevance.

Evaluating these configurations against the baseline (CNN+smax) revealed that substituting the temporal CNN with Temporal Attention (attn+smax) yielded comparable classification accuracies. This comparable performance corroborates the findings from the temporal shuffling experiments (**Supplementary Note 1**), confirming that the 1D-

CNN does not memorize rigid chronological sequences but operates as a dynamic weighting filter, conceptually analogous to an attention mechanism. Conversely, substituting the Softmax pooling with a Sigmoid activation (CNN+sigm, attn+sigm) induced a sensible performance degradation and increased cross-validation variance, particularly in the PDZ3 domain. The data indicates that while the specific temporal integration formalism is highly adaptable, enforcing a competitive feature selection mechanism (Softmax) constitutes the most robust configuration for isolating state-defining invariants across diverse physical systems.

Further evaluations concerning node features, embedding capacities, and spatial topologies yielded consistent behavior:

- *Dihedral Features:* The inclusion of continuous angular features, either restricted to the backbone ($\phi$, $\psi$) or expanded to the full side-chain ($\chi_1$, $\chi_2$), provided no significant performance improvement over the baseline, which relies exclusively on continuous inter-residue distances (RBFs) and biochemical node identities.

- *Embedding Dimensions:* Node embeddings define the representational capacity of the spatial message-passing filters for local residue environments, whereas global embeddings dictate the dimensionality of the final latent space utilized for contrastive learning and state classification. Scaling these latent dimensions down (32 global / 64 node) from the baseline configuration (64 global / 128 node) resulted in a slight performance degradation. Conversely, increasing the capacity (128 global / 256 node) yielded a marginal improvement in validation accuracy. However, this minor gain was accompanied by a substantial increase in computational overhead, specifically concerning GPU VRAM consumption during the training phase. Consequently, the baseline configuration was retained, as it provides the optimal balance between representational expressivity and computational tractability.

- *Spatial Cut-off:* Modulating the physical interaction cut-off demonstrated predictable topological effects. Reducing the cut-off to 8.0 Å caused a systematic drop in accuracy due to the loss of long-range spatial context. Expanding the cut-off to 12.0 Å marginally increased performance in the T4-Lysozyme system but introduced computational overhead and potential topological noise. The 10.0 Å threshold was confirmed as a practical balance between spatial resolution and computational efficiency.

**Temporal Sampling Parameters:** The stability of the spatiotemporal architecture is strictly dependent on the volume and resolution of the temporal data processed. To quantify this dependency, a discrete hyperparameter search was executed across varying combinations of stride and offset, utilizing a fixed window size of 25 frames (**Supplementary Fig. 3**). The analysis was conducted on the T4-Lysozyme, the PDZ3 Domain (non-masked) and the A2A receptor (agonist-bound vs apo non masked) systems. The offset parameter dictates the interval between the starting frames of consecutive sliding windows, determining the total number of windows extracted from a continuous trajectory. Decreasing the offset (e.g., from 25 to 1) maximizes the temporal overlap, substantially expanding the training dataset volume and generally improving classification accuracy. The stride parameter defines the step size utilized when sampling the discrete frames *within* a defined temporal window. Increasing the stride (e.g., from 1 to 20) effectively lowers the temporal resolution of the extracted dynamics and simultaneously restricts the mathematical boundary for window extraction. The heatmaps demonstrate that high stride values consistently degrade model performance.

Crucially, extreme parameter combinations (characterized by high offset and high stride) severely limit the absolute number of temporal windows available for gradient descent. In these data-starved regimes, the network fails to converge, and the classification accuracy collapses toward the random guess threshold (50.0%). The analysis confirms that minimizing both stride and offset maximizes validation accuracy; however, this scaling directly increases the computational cost of the training phase. The final baseline hyperparameters were subsequently selected to strictly operate within the converged, high-accuracy plateau while maintaining a practical balance with computational tractability (stride 1 offset 10).

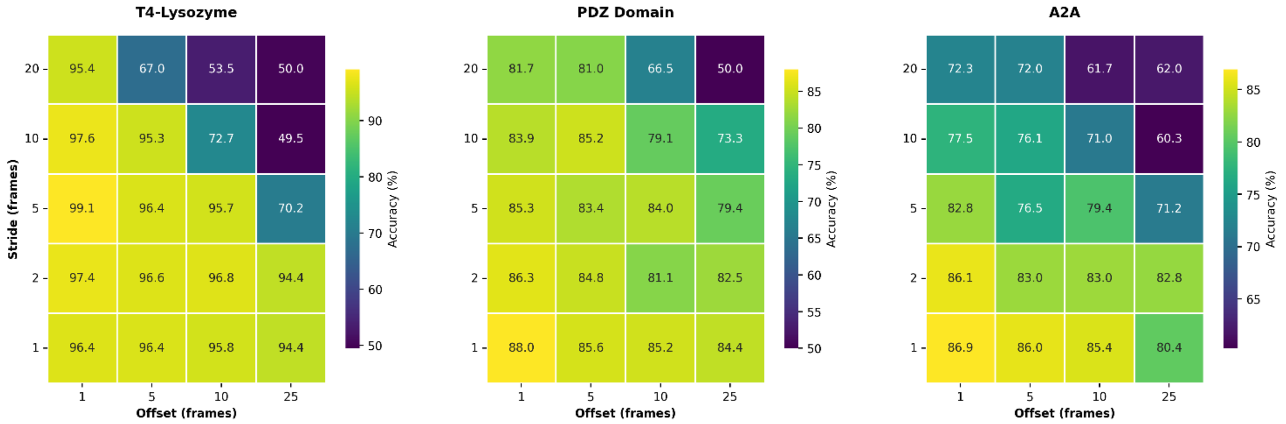


**Supplementary Fig. 5:** Sensitivity analysis of temporal sampling parameters (stride and offset). Evaluation of model performance across the hyperparameter space defining the temporal input construction. Discrete heatmaps displaying the exact mean validation accuracy (%) for each stride-offset combination. All experiments were conducted with a fixed temporal window size of 25 frames. The color gradient (purple to yellow) represents increasing classification accuracy.
